# Summary statistic analyses can mistake confounding bias for heritability

**DOI:** 10.1101/532069

**Authors:** John B. Holmes, Doug Speed, David J. Balding

## Abstract

LD SCore regression (LDSC) has become a popular approach to estimate confounding bias, heritability and genetic correlation using only genome wide association study (GWAS) test statistics. SumHer is a newly-introduced alternative with similar aims. We show using theory and simulations that both approaches fail to adequately account for confounding bias, even when the assumed heritability model is correct. Consequently, these methods may estimate heritability poorly if there was inadequate adjustment for confounding in the original GWAS analysis. We also show that choice of summary statistic for use in LDSC or SumHer can have a large impact on resulting inferences. Further, covariate adjustments in the original GWAS can alter the target of heritability estimation, which can be problematic for test statistics from a meta-analysis of GWAS with different covariate adjustments.

## Introduction

LD SCore regression (LDSC) uses genome-wide association test statistics to estimate confounding bias, the heritability tagged by SNPs 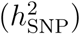, how 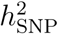 is distributed across the genome and the genetic correlation of pairs of traits (Bulik-Sullivan et al., 2015a,b; Finucane et al., 2015; Gazal et al., 2017). Its use of test statistics rather than individual genotype data means that it is effectively unlimited in sample size, and can make use of published studies that do not release the genotypes of participants. Moreover the test statistics can be obtained from a single GWAS or from a meta-analysis of multiple GWAS. These advantages have led to LDSC being very widely used.

LDSC regresses the test statistic at each SNP on an “LD score”, defined as a sum of linkage disequilibrium (LD) coefficients over neighbouring SNPs. The regression slope and intercept are interpreted as, respectively, 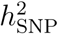 and confounding bias not corrected in the GWAS analysis. SumHer (Speed and Balding, 2019) and S-LDSC (Finucane et al., 2015; Gazal et al., 2017) generalise LDSC by introducing weights into the LD score. The weights correspond to a heritability model that relates the expected heritability of a SNP to its properties known *a priori*. SumHer uses fixed, SNP-specific weights reflecting LD and minor allele fraction (MAF). In the most recent version of S-LDSC (Gazal et al., 2017), weights based on LD and MAF as well as functional annotations are estimated in the summary statistic analysis. HESS (Shi et al., 2016) and RSS (Zhu and Stephens, 2017) are other summary statistic methods that require more information than association test statistics.

Researchers using LDSC or SumHer usually have not performed the underlying GWAS analyses, but use test statistics obtained from public data repositories (Zheng et al., 2017) that may lack information needed to check the assumptions underlying these methods. Here, we examine the validity of these assumptions under a range of scenarios. We do not revisit the topic of the underlying heritability model (Speed and Balding, 2019), rather we will highlight problems that arise even when the simulation and analysis heritability models are the same.

We derive expected values of association statistics and show that confounding effects are SNP dependent, and correlated with LD score (Appendices 2-3), contravening a fundamental assumption of LDSC and SumHer. Thus a global adjustment term can fail to remove confounding effects, although a multiplicative adjustment can correct over-conservatism arising from use of genomic control (Speed and Balding, 2019).

We illustrate the magnitude of the problem through simulations. Our investigation covers two summary statistics and we show that inferences from LDSC or SumHer can be greatly impacted by this choice. Further, we show that the definition of 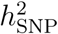 targeted by LDSC and SumHer varies with the covariates fitted in the GWAS analyses. This can be important in meta-analysis: a summary statistic heritability analysis based on studies with different covariate adjustments will merge estimates of different parameters.

### A general model

We derive approximations for E[*S*_*j*_] and perform simulation studies using a highly general phenotype model,

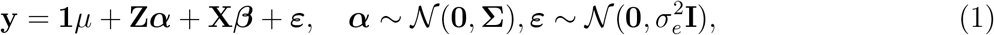

where *µ* is an intercept, **Z** an *n* × *m* matrix of standardised SNP genotypes, ***α*** a vector of SNP effect sizes, and **Σ** a diagonal matrix with *j*th entry 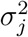. The *n* × *p* matrix **X** contains column-standardised covariate values, while ***β*** is a vector of covariate effects. If Cor(**X**, **Z**) ≠ **0**, the **X** are confounders for genetic association analysis. The most important example is population structure when both **X** and **Z** vary with, for example, geography or social strata.

The GCTA model (Yang et al., 2011a) is the special case of (1) in which 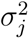 is constant over SNPs. This assumption also underlies LDSC. The LDAK model (Speed et al., 2017) is another special case of (1) in which 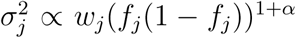, where the *w*_*j*_ reflect local LD at SNP *j*, *f*_*j*_ is the MAF of SNP *j* and *α* is a parameter that reflects selection on the phenotype. The value *α* = −0.25 has been shown to fit well over many human phenotypes (Speed et al., 2017). SumHer can be used with any heritability model, but here we implement SumHer with the LDAK values for 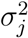 and *α* = −0.25.

When (1) is used as a simulation model, usually only a subset of the available SNPs are assigned non-zero effects. When used as an analysis model, because the causal SNPs are unknown all available SNPs should be included in (1). This mismatch between simulation and analysis models arises as it is impossible in practice to limit analyses to causal SNPs.

### Choice of test statistics

LDSC and SumHer both fit a linear regression to summary statistics *S*_*j*_, *j* = 1, …, *m*, obtained from a GWAS on *n* individuals:

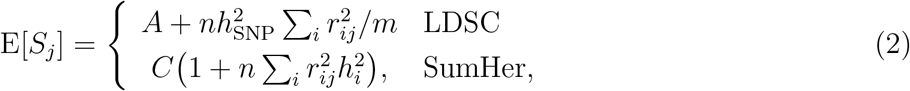

where 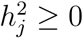 is interpreted as the expected heritability attributable uniquely to SNP *j*, with 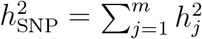. In (2), 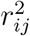 is an estimated LD coefficient with *r*_*ii*_ = 1 (see Methods), while *A* and *C* are alternative adjustments for confounding effects not accounted for in the GWAS analysis. Estimates of *C* using SumHer were reported to be much lower than the corresponding estimates of *A* from LDSC (Speed and Balding, 2019), but this was due to the difference in heritability model rather than whether the confounding term was additive or multiplicative. SumHer found that many GWAS had over-corrected for confounding (*C* < 1), whereas LDSC analyses of the same data typically found *A* > 1, indicating a need for further confounding adjustment (Speed and Balding, 2019).

In practice, *S*_*j*_ is often the Wald statistic 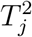 from a classical simple linear regression (Yang et al., 2011b; Bulik-Sullivan et al., 2015a; Finucane et al., 2015), which can be inferred from *p*-values. Its null distribution is *F*_1,*n*−2_, which converges to 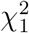 as *n* increases. However, LDSC was proposed assuming that 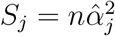, where *α*_*j*_ is the effect of SNP *j* when both **Z** and **y** are standardised (Bulik-Sullivan et al., 2015a). Assuming that no covariates were fitted, *S*_*j*_ is *n*/(*n*−1) times the standardised regression sum of squares 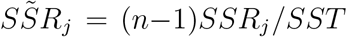. When covariates are included this equality no longer holds, but we check using simulation that 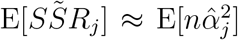, and where convenient we compute 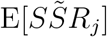 rather than 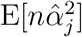.

Recently, GWAS test statistics have often been derived from a mixed regression model (Lippert et al., 2011; Loh et al., 2015) in which SNP *j* is tested while other SNPs are used to compute the variance structure of a random effect modelling cryptic kinship and population structure. We report the expectation of a general mixed model test statistic, finding that it depends on quantities not usually available from GWAS data repositories and that in general 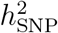 is non-identifiable (Appendix 3.7).

### Two definitions of 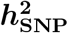

We define 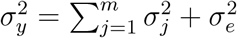, the phenotypic variance after conditioning on covariates/confounders **X**. However **X** may be unrecorded, or omitted from analysis, in which case 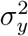 cannot be estimated, and only the total phenotypic variance 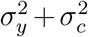 is available, where 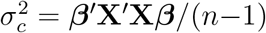 and *′* denotes transpose. This leads to two definitions of the heritability of SNP *j* (Weissbrod et al., 2018):

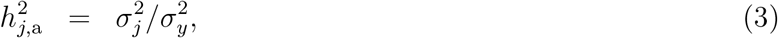

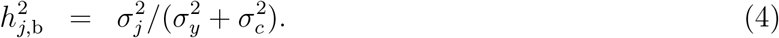

The conditional heritability, 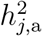, is standard when the **X** are modelled as fixed effects (Mrode, 2014; Pirinen et al., 2013), while the marginal heritability, 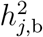, is usually preferred for random-effect covariates (Heckerman et al., 2016; De Villemereuil et al., 2018). We use 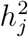 when there are no covariates or it is unimportant to distinguish 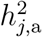 from 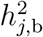. Henceforth we assume that the phenotype vector **y** is sample standardised, in which case 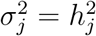.

## Methods

### Data processing

We used genotypes from the eMERGE network (Verma et al., 2014), following the same quality control steps as Speed and Balding (2019). From the 25,875 individuals, we randomly selected 8000 to form the study population, simulated their phenotypes and computed GWAS summary statistics. The remaining 17,875 individuals were used as a reference panel to compute *r*^2^ values for the summary statistic analyses.

We also generated three meta-analyses by dividing the study population randomly into two studies of size 4000, and calculating summary statistics for each study, both without and with covariate adjustment. Each meta-analysis used within-study phenotype standardisation, and computed 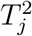 using inverse-variance weighting (Willer et al., 2010).

Of the SNPs remaining after quality control, 558,431 had non-zero LDAK weights (Speed et al., 2012) and only these SNPs contribute causal effects under the LDAK model and to SumHer analyses. We also restricted LDSC analyses, and simulations under the GCTA model (Yang et al., 2011a), to a set of 558,431 randomly-chosen SNPs.

### Simulation of phenotypes and summary statistics

For 150 iterations, we randomly sampled 35,000 causal SNPs and, under each of the GCTA and LDAK models, we generated five phenotypes with different covariate and confounding effects such that 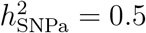 in all cases: **y**_*A*_ (no covariates or confounding), **y**_*B*_ (covariate effect, no confounders), and **y**_*Ci*_, *i* = 1, 2, 3 (confounding). For **y**_*B*_, **X** has two columns, and the simulated effects were such that 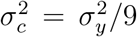, so that 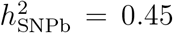. To explore incomplete control of confounding, for all **y**_*C*_ phenotypes confounders correspond to a two-level hierarchical population structure. First, three subpopulations were identified using *k*-means clustering on the leading two principal components (PCs) of the SNP correlation matrix **Z**^*T*^**Z**/*m*, restricted to SNPs with non-zero LDAK weight in order to minimise any effect of correlated SNPS.

Within each of these subpopulations, three sub-subpopulations were defined by *k*-means clustering on the two leading PCs computed only from subpopulation members. We assigned different phenotype means to the nine sub-subpopulations, while SNP effect sizes remained the same. For **y**_*C*_ phenotypes we consider both **X** corresponding to the three subpopulations (two columns), and **X** corresponding to all nine sub-subpopulations (eight columns). The 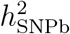 values were 0.45 (C1), 0.475 (C2) and 0.49 (C3), corresponding to, respectively, 10%, 5% and 2% of the phenotypic variation due to confounding. **y**_*A*_ phenotypes and principal components were calculated using the LDAK software, while *k*-means clustering and the simulation of **y**_*B*_ and **y**_*C*_ phenotypes was undertaken in R (R Core Team, 2016).

For all phenotypes we compute 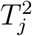 and 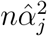, both with and without adjusting for covariates **X**. Based on these statistics we estimate 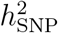 using SumHer for LDAK phenotypes (results in main text) while for GCTA phenotypes (Appendix 4.1) we used LDSC as implemented in the LDAK software (Speed et al., 2012). The two sets of results are broadly similar; we comment in the text on notable differences.

### Large-effect SNPs

In part because of the problem of the unknown number of causal SNPs, but also due to model mis-specification such as incomplete control of confounding, in many GWAS values of *S*_*j*_ arise that are extreme outliers under the assumed analysis model. Ideally, the solution would be to improve the analysis model, for example using a distribution with thicker tails than the Gaussian, or assigning an atom of prior probability at each SNP to a zero effect. However, because of computational advantages associated with model (1), in practice an ad-hoc data filtering approach is often adopted in which a SNP is removed if its estimated effect size is too large to be well-supported under the model. As we have control over confounding in our simulations, our main results do not use filtering. In Appendix 4.2, we consider the impact of filtering, where we follow Zheng et al. (2017) and exclude from analysis any SNP with *S*_*j*_ > 80. In the analysis of **y**_*A*_ and **y**_*B*_ simulations, no SNP was excluded, while for 32 C1, 2 C2, and 0 C3 LDAK and 44 C1, 14 C2, and 0 C3 GCTA simulations, at least one SNP was excluded for both 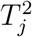 and 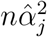 when **X** was ignored in the GWAS analysis.

## Results

For derivations of the expectations given below, see Appendix 3. Simulation results reported here are for SumHer analyses of LDAK phenotypes (see Methods); corresponding results using LDSC analyses of GCTA phenotypes are broadly similar (Appendix, Figures S1-4).

### No confounding (Cor(X, Z) = 0)

For a single GWAS with no covariate/confounder effects

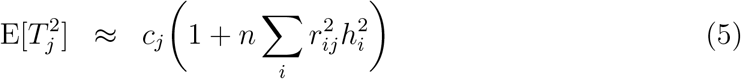

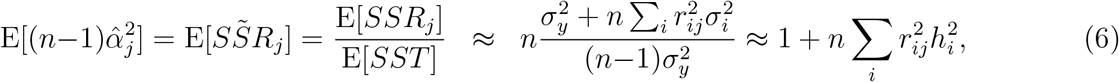

where 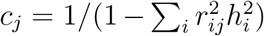. For complex traits 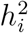 is typically small, so that *c*_*j*_ slightly exceeds 1 for many *j*. Both (5) and (6) are special cases of the SumHer model (2), but the deviation of *C* from unity in (5) is not due to confounding.

SumHer estimates of 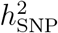 based on GWAS summary statistics in the absence of covariate/confounder effects are centred close to the true value of 0.5 for both statistics (Figure 1(a)), so that for our simulations the deviation of the *c*_*j*_ from 1 appears to be negligible. The mean estimate of 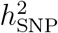 does not noticeably change when *A* or *C* is estimated rather than fixed at the true values (*A* = *C* = 1), but the variance increases due to uncertainty arising from the additional parameter estimation.

**Figure 1:**
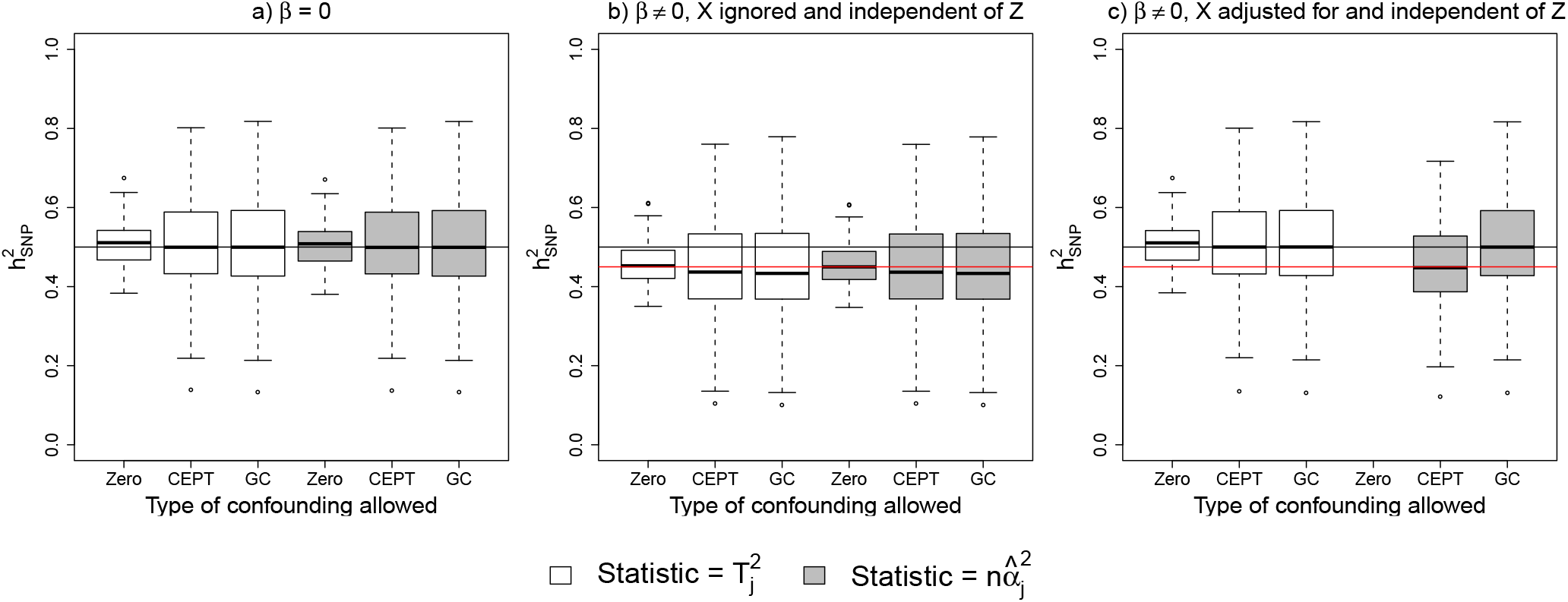
Estimates of 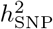 obtained by analysis of summary statistics from simulated GWAS. The black and red horizontal lines indicate the values of 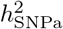 and 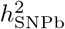, the SNP heritability without and with conditioning on covariates. Zero, CEPT and GC refer to no, *A* and *C* confounding terms being estimated in the analysis. Phenotypes are simulated under the LDAK model and the analyses performed using SumHer. (a) Phenotypes with no covariate effects. (b) Phenotypes with covariate effects but **X** ignored in the analysis. (c) Phenotypes with covariate effects and **X** adjusted for in the analysis. The “Zero” estimates when 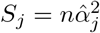 are all negative and are not shown.

When covariates affect **y** but **X** is ignored in the GWAS analysis, 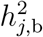 is estimated rather than 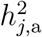 because now 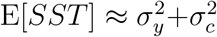 rather than 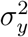. Again, the average estimate of 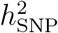 changes little when *A* or *C* is estimated rather than fixed at 1 (Figure 1(b)). When **X** is included in the GWAS analysis,

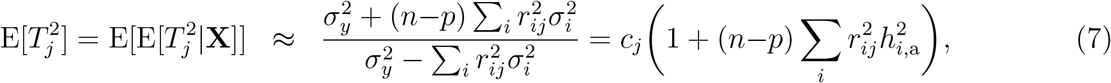

which is the same as (5) but with *n*−*p* in place of *n* and 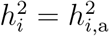. Further,

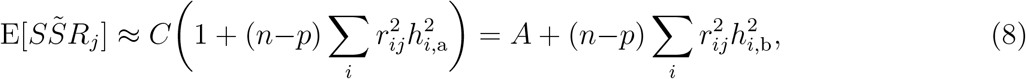

where 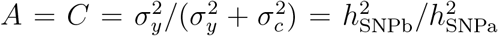. Figure 1(c) reflects properties evident from (7) and (8): only 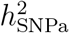 can be estimated from 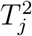, whereas either 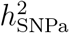 or 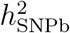 can be estimated from 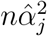, according to whether *C* or *A* is fitted.

Figure 2 shows that 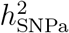 and 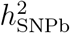 are estimated with no apparent bias if, respectively, both studies did and did not adjust for covariates in a two-GWAS meta-analysis. Again, the inclusion of *A* or *C* terms has little effect on the mean estimates, as there is no confounding. When there is a mismatch in covariate adjustments between the two GWAS, the estimate of 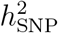 is intermediate between 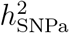 and 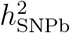 (Figure 2, green bars). In practice many meta-analyses do combine studies with different covariate adjustments, which may not adversely affect association tests but does affect heritability analyses. Examples include the meta-analyses of height (Lango et al., 2010) and blood pressure (The International Consortium for Blood Pressure Genome-Wide Association Studies et al., 2011) re-analysed using LDSC (Bulik-Sullivan et al., 2015a), and those of psychiatric traits (Okbay et al., 2016) and Type 2 diabetes (Scott et al., 2017).

**Figure 2:**
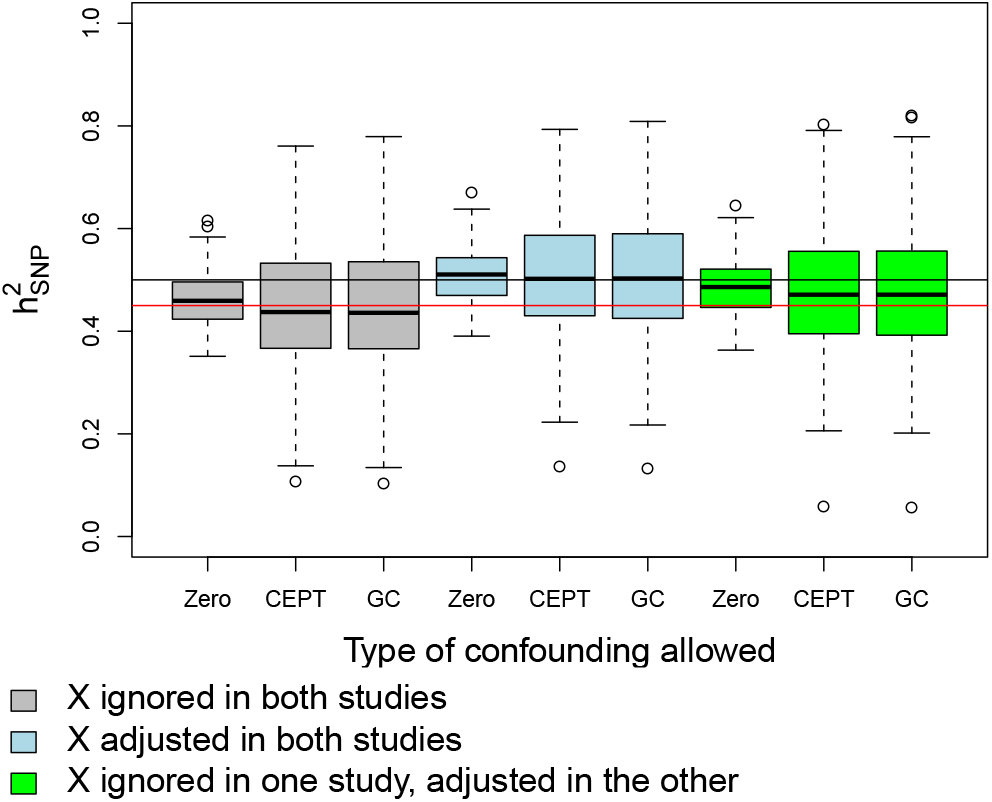
Estimates of 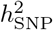 obtained from SumHer analysis of summary statistics calculated from a meta-analysis of two GWAS. The black and red horizontal lines indicate the values of 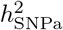 and 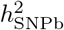. Zero, CEPT and GC refer to no, *A* and *C* confounding terms in the analysis model.

### Confounding (Cor(X, Z) ≠ 0)

When confounder **X** is ignored in the GWAS analysis:

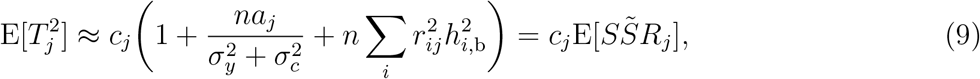

where 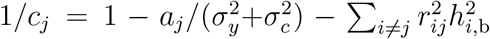 and 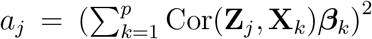 with **X**_*k*_ denoting column *k* of **X**. Assuming *c*_*j*_ ≈ 1, only 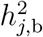 is estimable, and (9) includes an additive constant resembling *A* in (2). However, this term is SNP-dependent, and for it to correspond to *A* in (2) we require *a*_*j*_ to be independent of LD score, which typically does not hold (see Appendix 2). Instead, we expect the estimate of 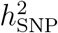 to be inflated by an amount that depends on both Cov(**X**, **Z**) and 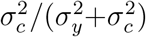. Replacing (9) with the SumHer regression model leads to similar difficulties.

When **X** is included in the GWAS analysis, the estimated SNP effect, 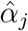, can be obtained from the linear regression of the residuals of **y**|**X** on the residuals of **Z**|**X** (Frisch and Waugh, 1933), and

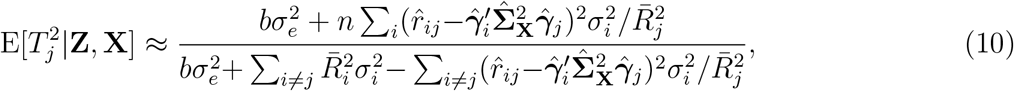

where 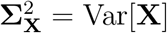 and, from the regression of **Z**_*j*_ on **X**, 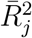 is one minus the coefficient of determination and 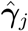 is the vector of estimated coefficients, while *b* = (*n*−p−2)/(*n*−2). Further,

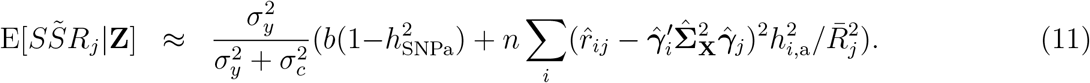

Now, 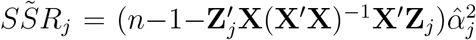 and, unlike when Cor(**X**, **Z**) = **0**, the term multiplying 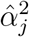 varies over SNPs.

As expected, ignoring confounders results in inflated estimates of 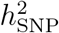 (Figures 3(a) and 4(a, b)). The values of 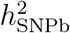, are 0.45 for *C*1, 0.475 for *C*2, and 0.49 for *C*3 phenotypes, yet the average estimates are in the reverse order (*C*1 > *C*2 > *C*3) because of the inadequately-corrected confounding (Figure 4(a,b)).

**Figure 3:**
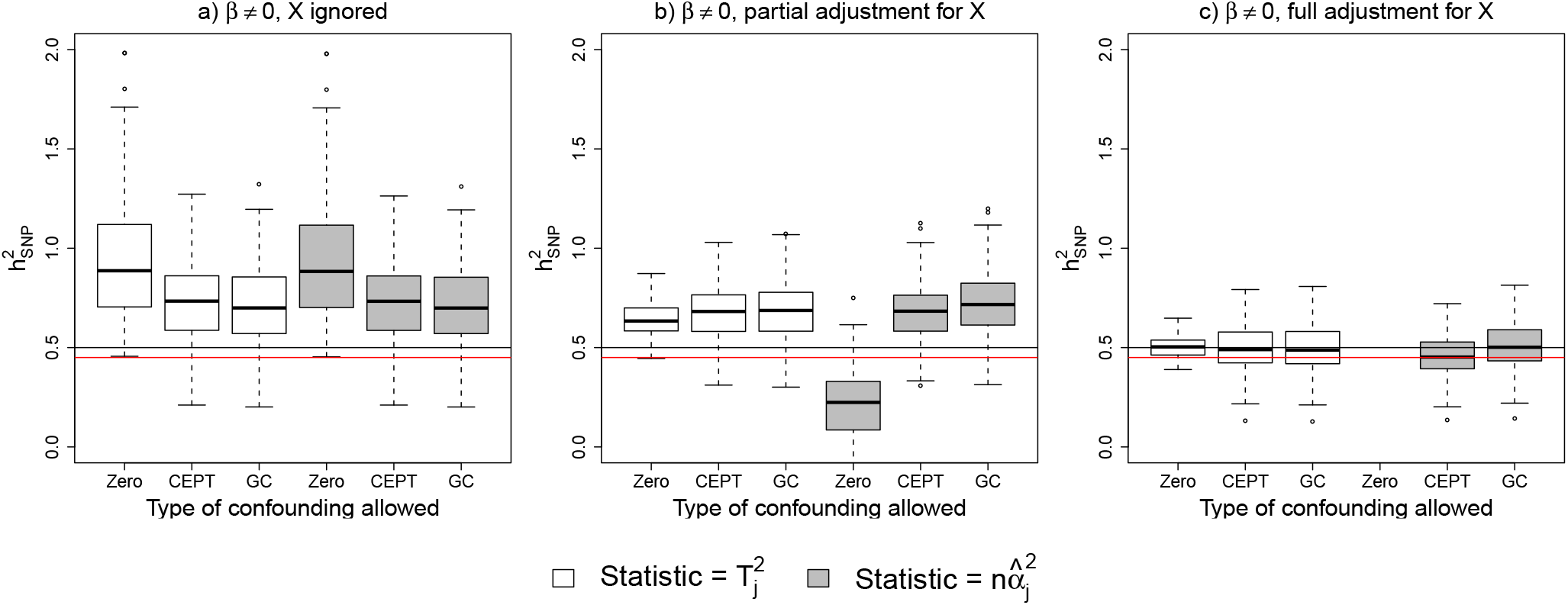
Similar to Figure 1, but here GWAS phenotypes are subject to confounding: phenotype means differ among three subpopulations that each consist of three sub-subpopulations. Subpopulations were constructed by applying k-means clustering to principal components of the SNPs with non-zero LDAK weight. Estimates of 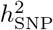 from a GWAS with (a) no covariate adjustment, (b) adjustment for the three subpopulations but not the sub-subpopulations, (c) full covariate adjustment.

**Figure 4:**
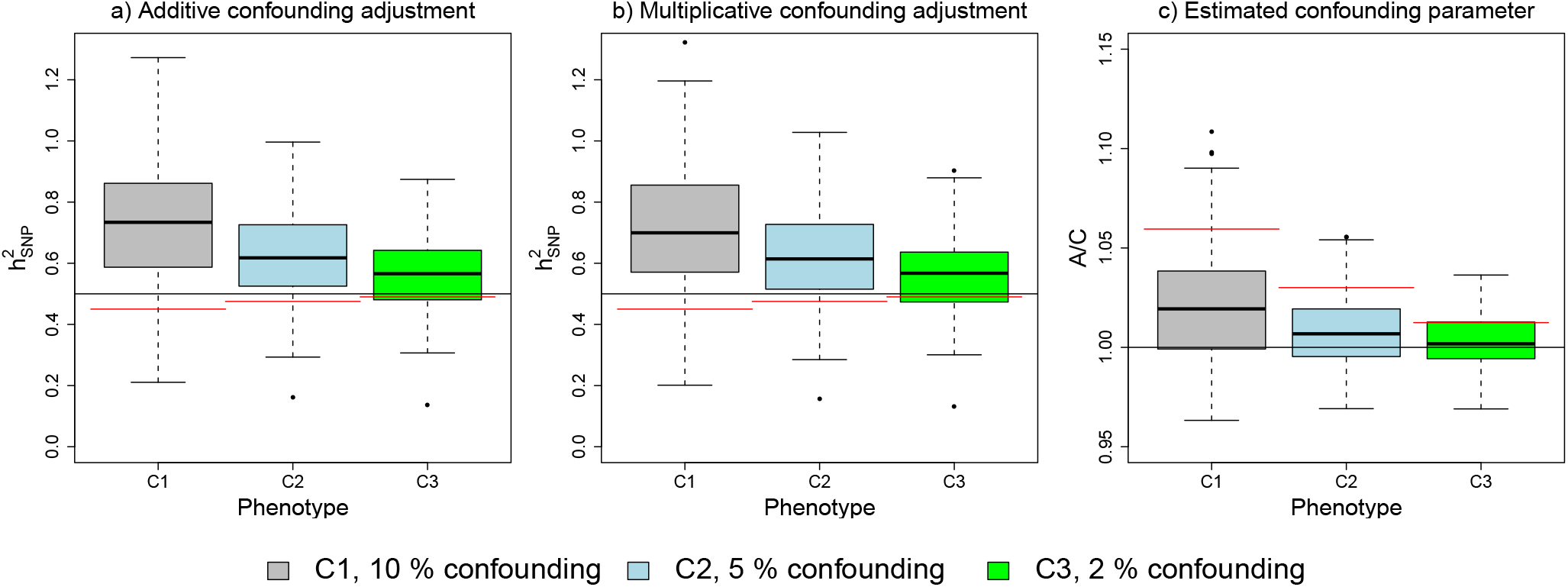
Estimating 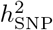 and confounding parameters from phenotypes when 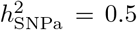 and 10% (C1), 5% (C2) and 2% (C3) of phenotypic variance due to confounding. See Figure 3 for details of the confounding. The black lines in (a,b) indicate the simulated value of 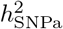 and the red lines the simulated value of 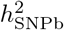, while the box plot shows the distribution of estimates when applying the confounding adjustment indicated in the plot heading. In (c), the black line corresponds to *A/C* = 1 corresponds to zero confounding bias and the red line the mean level of confounding, estimated as 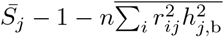. Note that the *y*-axis differs between (a,b) and (c).

The *A*/*C* estimates are consistently too low (Figure 4(c)) because the positive association between *a*_*j*_ and LD score leads to some of the confounding being misinterpreted as heritability. Comparing *C*1, *C*2 and *C*3 phenotypes, we find that the bias in 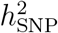 is, like *a*_*j*_, a function of 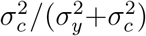, the proportion of phenotypic variance due to confounding. Figure 4(c) shows average *A*/*C* estimates > 1 in the presence of confounding, but this does not always hold, and estimates *A* < 1 have been reported, such as in GWAS of rheumatoid arthritis (Bulik-Sullivan et al., 2015a), age at first birth and number of children (Barban et al., 2016), body mass index and hip-waist ratio (Speed and Balding, 2019), which could be due to confounding of the type considered here. The level of bias in 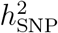 was higher in the LDAK simulations than for the GCTA simulations (Appendix 4.1).

Our finding of inadequate adjustment for confounding is concordant with the results of two recent analyses of stratified populations (DeVlaming et al., 2017; Berg et al., 2018), but not with Lee et al. (2018) who considered confounding by parental genotype. This is because parental average genotype generates an additive genetic effect, which inflates the slope but not the intercept of the summary statistic regression.

Partial covariate adjustment in the GWAS analysis reduced but did not eliminate bias in 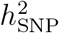 estimates (Figure 3(b)) and led to divergence of the estimates based on 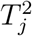 and 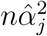. Full covariate adjustment did lead to unbiased estimates of 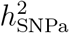 when 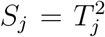 whether *A* or *C* or neither was fitted (Figure 3(c)). When 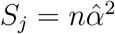, fitting *A* or *C* led to estimates of 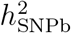 and 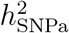, respectively. These results indicate that although confounding adjustment can mask some of the heritability signal, which is intuitively why (10) and (11) differ from (7) and (8), the differences appear negligible in this case. However, for populations with much stronger stratification adjusted for in the GWAS stage, the distinction between (10), (11) and (7), (8) was reported to be important for estimating 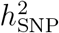 (Luo et al., 2018).

### Sample overlap and mis-specification of the heritability model

Our focus has been on heritability analyses when the underlying GWAS failed to account for all covariates or confounders. Here we briefly discuss some other possible issues that complicate summary statistic heritability analysis.

First, consider a meta-analysis in which some individuals have been included in multiple studies, but all non-genetic effects were accounted for in the component studies. In this case, the expected test statistic will be additively inflated (see Appendix 3.6 and Figure 5), and estimation of 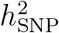 remains unbiased. Furthermore, if the level of overlap is known, then the inflation in the expected statistic can be predicted in advance. In contrast, the population structure in our simulations implies some relatedness among all individuals, leading to a relationship between confounding and LD that cannot be modelled using *A* or *C*. The example of non-trivial intercepts in Lee et al. (2018) appropriately correcting for confounding is similar to sample overlap, as it was based on twins, so can be viewed as two dependent samples, each consisting of unrelated individuals.

**Figure 5:**
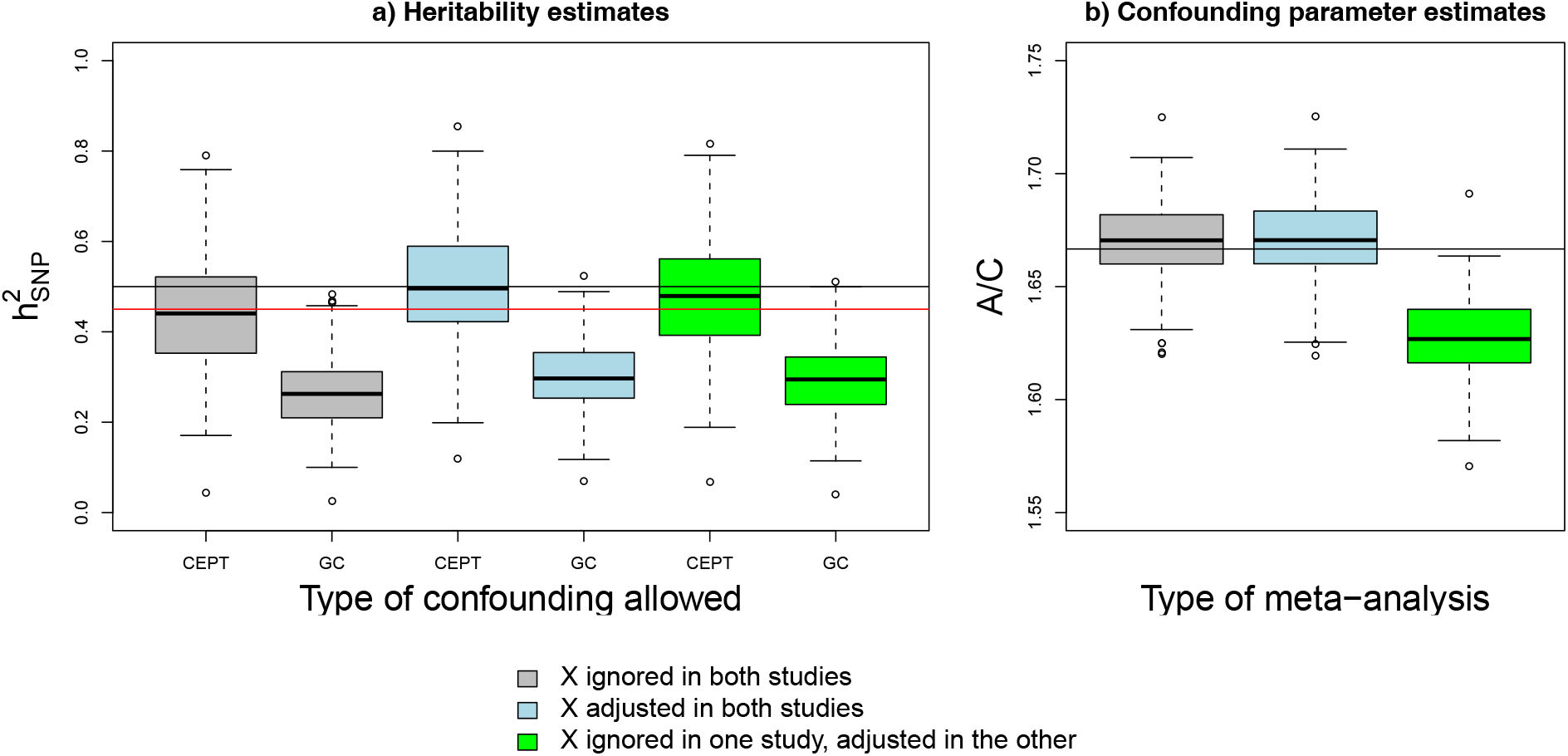
Similar to Figure 2, but now the meta-analysis includes overlapping 4000 individuals out of a total of 12,000. The black and red horizontal lines in (a,c) indicate the values of 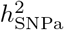 and 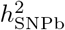. The black horizontal line in (b,d) indicates the expected intercept. CEPT and GC refer to A and C confounding terms in the analysis. Plot (a) gives estimates of 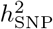 obtained under either an A or C view of the confounding parameter, while plot (b) gives estimates of the confounding parameter. Phenotypes were simulated in accordance to a LDAK model.

Figure 6 (a) shows that use of a well-chosen reference panel (which is the case for our simulations) has minimal impact on parameter estimates compared with computing LD coefficients from the GWAS genotypes (In-sample estimates). In Figure 6 (b-d), we compare the relationship of the predictor 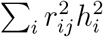 with summary statistics, under some extreme scenarios to illustrate sensitivity to mis-specification of the heritability model. If the true heritability model was LDAK, but the summary statistic analysis assumed a GCTA model, 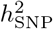 would be under-estimated, and the confounding parameter *A* over-estimated (Figure 6 (b), see also Speed and Balding (2019)). If all causal variants have low MAF (Figure 6 (c)), then 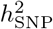 is again under-estimated. If causal variants are all in regions of high LD (Figure 6 (d)), then 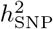 is over-estimated and *A* is under-estimated.

**Figure 6:**
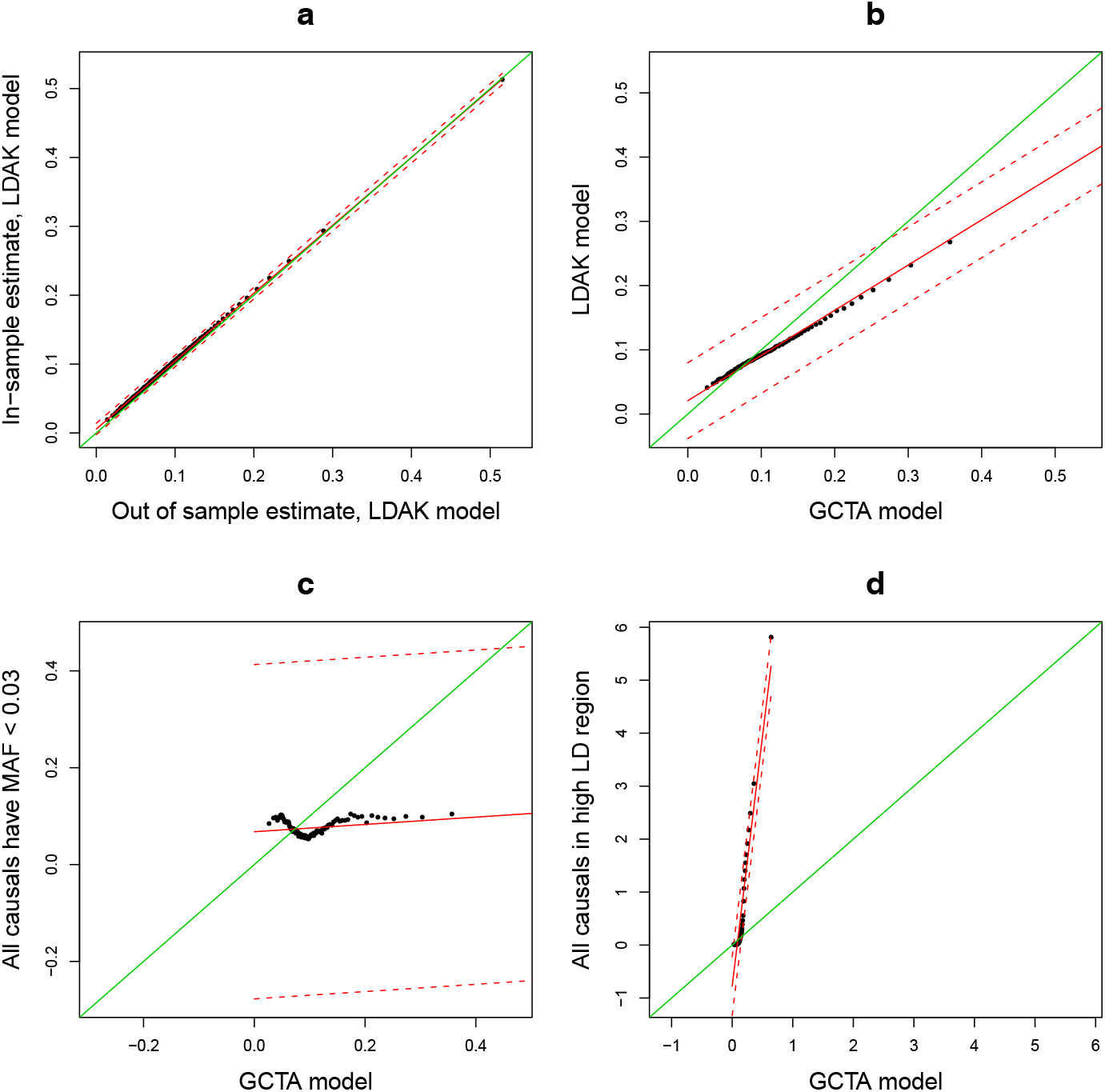
Comparing the slope of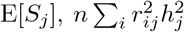, for a range of heritability models. In (a), the slopes given by assuming a LDAK model where 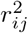 is estimated by an external reference panel (x-axis) and using sample genotypes (y-axis) are compared. (b) compares the slopes assuming a GCTA (x-axis) and LDAK (y-axis) model. (c) compares the slopes assuming a GCTA model (x-axis) and a heritability model where only variants with MAF < 0.03 are causal (y-axis). (d) compares the slopes assuming a GCTA (x-axis) and a heritability model where only SNPs where 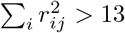 are causal (y-axis). In (c) and (d), causal variant effects are assumed 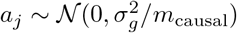. where *m*_causal_ is the number of causal variants. The green line in the plots is the 45 degree line, while the red lines correspond to the the line of best fit (solid) and associated 95 % prediction intervals. The dots are percentile bins.

## Discussion

We have shown theoretically, and illustrated using simulation, that GWAS confounding bias is in general SNP dependent and correlated with LD. Therefore if the original GWAS analysis did not avoid confounding effects, heritability analysis of GWAS summary statistics using regression on an LD-related predictor, as implemented in LDSC (Bulik-Sullivan et al., 2015a) and SumHer (Speed and Balding, 2019), can fail to adequately account for confounding bias, leading to poor 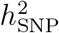 estimation. To avoid this, we propose that such analysis proceeds in practice by first fitting a model with *A* or *C* unconstrained, and testing for deviation of its value from one. If *A/C* is significantly different from one subsequent heritability analysis would be unreliable. By this criterion, heritability analysis should proceed for only 7 out of 24 LDSC analysis of GWAS reported in Table 1 Bulik-Sullivan et al. (2015a), and 3/25 LDSC analyses and 17/25 SumHer analyses of GWAS reported in Supplementary Table 2 of Speed and Balding (2019).

When covariates were fitted in the GWAS analysis, we found very different results according to whether *S*_*j*_ was chosen to be the Wald statistic 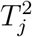 or the statistic 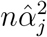 used to justify LDSC (Bulik-Sullivan et al., 2015a). Further, the estimable definition of 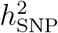 varies with the covariate adjustment performed in the original association analysis. The statistic 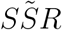, closely related to 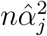, can be used to estimate 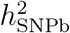 regardless of the (non-confounder) covariates fitted, and hence a valid meta-analysis of 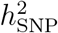 estimates is possible. However, 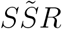 is often not available in published GWAS results, and like 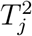 it is subject to SNP-dependent confounding that can bias estimates of 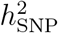. We have also shown that mis-specification of the heritability model can cause a regression intercept different from one, and biased 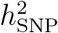estimates, even if the underlying GWAS correctly accounted for confounders.

One source of error in published GWAS test statistics is genomic control, which applies a common multiplicative adjustment, derived under an assumption of sparse causal effects. It tends to over-adjust for highly-polygenic traits, which is corrected using SumHer (Speed and Balding, 2019).

We have only considered quantitative phenotypes, and we have not examined in detail the question of the validity of 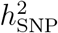 analyses based on mixed model association statistics. In the appendix, we show that the exact expected value of such statistics contain terms not usually obtainable from public databases. Further 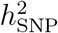 is non-identifiable without further assumptions, which if ignored could lead to severely biased estimates.

Our findings accord with those of others (DeVlaming et al., 2017; Berg et al., 2018), and some statements/results in the original LDSC paper (Bulik-Sullivan et al., 2015a). For example, Bulik-Sullivan et al. (2015a) found that some heritability was inferred in simulations of confounding-only phenotypes, which was attributed to linked selection generating the correlation between confounding effect and LD score. Further, their interpretation of the intercept was based on average results from distinct populations, not replicate samples from the same population. This ignores the structure, and hence confounding, that is specific to a population. Supplementary Table 4C in Bulik-Sullivan et al. (2015a) shows that in the presence of confounding only 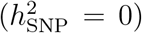, one population (cou3) has higher LDSC 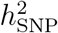 estimates for all three traits (0.144, 0.254, 0.229) than for any of the other 18 trait/population combinations (average: 0.030). Most importantly, the claim that LD score is not associated with confounding (Bulik-Sullivan et al., 2015a) was based on marginalising over the confounding term *a*_*j*_, which is inappropriate as *a*_*j*_ and LD score are both SNP specific.

## Acknowledgements

DS is funded by the European Unions Horizon 2020 Research and Innovation Programme under the Marie Sk-łodowska-Curie grant agreement number 754513, by Aarhus University Research Foundation (AUFF) and the Independent Research Fund Denmark under Project 7025-00094B. DB is funded by the Australian Research Council under grant DP190103188. The eMERGE Network was initiated and funded by NHGRI through the following grants: U01HG006828 (Cincinnati Childrens Hospital Medical Center/Boston Childrens Hospital); U01HG006830 (Childrens Hospital of Philadelphia); U01HG006389 (Essentia Institute of Rural Health, Marshfield Clinic Research Foundation and Pennsylvania State University); U01HG006382 (Geisinger Clinic); U01HG006375 (Group Health Cooperative); U01HG006379 (Mayo Clinic); U01HG006380 (Icahn School of Medicine at Mount Sinai); U01HG006388 (Northwestern University); U01HG006378 (Vanderbilt University Medical Center); and U01HG006385 (Vanderbilt University Medical Center serving as the Coordinating Center). Access to eMERGE Network data was granted under dbGaP Project 14422, ‘Comprehensive testing of SNP-based prediction models’.

## Declaration of interests

The author declare there are no conflicts of interest.

