## Appendix for "Summary statistic analyses can mistake confounding bias for heritability"

#### 1 The underlying model for our work

Throughout this paper, we assume that the phenotype model can be represented as,

$$\mathbf{y} = \mathbf{1}\mu + \mathbf{X}\boldsymbol{\beta} + \mathbf{Z}\boldsymbol{\alpha} + \boldsymbol{\varepsilon}, \quad \boldsymbol{\alpha} \sim \mathcal{N}(\mathbf{0}, \boldsymbol{\Sigma}), \boldsymbol{\varepsilon} \sim \mathcal{N}(\mathbf{0}, \sigma_e^2\mathbf{I}), \tag{1}$$

making no distributional assumption for $\boldsymbol{\beta}$ and assuming that columns of $\mathbf{X}, \mathbf{Z}$ are standardised. In particular we will focus on what impact any associations between $\mathbf{X}$ and $\mathbf{Z}$ have on expected association statistics, and the implied associations in quadratics of $\mathbf{X}, \mathbf{Z}$.

#### 2 Association between LD score and confounding

To simplify the presentation we only consider the LD score as defined in LDSC, and not the weighted version used by SumHer, although similar issues do arise for SumHer. We assume without loss of generality that $\mathbf{X}$ is orthonormal with $< n$ columns, and $\mathbf{Z}$ is standardised. Let $\mathbf{Z} = \mathbf{X}\mathbf{B} + \mathbf{E}$ where $\mathbf{B} \perp \mathbf{E}$. Then

$$\begin{aligned}
\mathrm{Cor}(\mathbf{Z}) &= \mathrm{Cov}(\mathbf{Z}) = \mathrm{Cov}(\mathbf{X}\mathbf{B} + \mathbf{E}) = \mathbf{B}'\mathrm{Cov}(\mathbf{X})\mathbf{B} + \mathrm{Cov}(\mathbf{E}) = \mathbf{B}'\mathbf{B} + \mathrm{Cov}(\mathbf{E}), && (2)\\
\mathrm{Cov}(\mathbf{Z}_j, \mathbf{X}\boldsymbol{\beta}) &= \mathrm{Cov}(\mathbf{X}\mathbf{B}_j, \mathbf{X}\boldsymbol{\beta}) = \mathbf{B}_j'\mathrm{Cov}(\mathbf{X})\boldsymbol{\beta} = \mathbf{B}_j'\boldsymbol{\beta}. && (3)
\end{aligned}$$

Hence $(\mathbf{B}_j'\boldsymbol{\beta})^2 = a_j$ and we will write $\mathbf{B}_j'\boldsymbol{\beta} = \sqrt{a_j}$ which can be positive or negative. Now decompose $\mathbf{B}_j$ into a multiple of $\boldsymbol{\beta}$ plus a vector $\hat{\boldsymbol{\epsilon}}_j$ that is orthogonal to $\boldsymbol{\beta}$. Then the coefficient of $\boldsymbol{\beta}$ is $\boldsymbol{\beta}'\mathbf{B}_j/\boldsymbol{\beta}'\boldsymbol{\beta} = \sqrt{a_j}/\boldsymbol{\beta}'\boldsymbol{\beta}$. Substituting (2) into the LD score expression, we obtain

$$\begin{aligned}
\sum_i r_{ij}^2 &= \sum_i \mathrm{Cor}(\mathbf{Z})_{ij}^2 = \sum_i (\mathbf{B}_i'\mathbf{B}_j + \mathrm{Cov}(\mathbf{E})_{ij})^2 = \sum_i ((\boldsymbol{\beta}\sqrt{a_i}/\boldsymbol{\beta}'\boldsymbol{\beta} + \hat{\boldsymbol{\epsilon}}_i)'(\boldsymbol{\beta}\sqrt{a_j}/\boldsymbol{\beta}'\boldsymbol{\beta} + \hat{\boldsymbol{\epsilon}}_j) + \mathrm{Cov}(\mathbf{E})_{ij})^2 \\
&= \sum_i (\sqrt{a_i a_j}/\boldsymbol{\beta}'\boldsymbol{\beta} + \hat{\boldsymbol{\epsilon}}_i'\hat{\boldsymbol{\epsilon}}_j + \mathrm{Cov}(\mathbf{E})_{ij})^2 \\
&= \frac{a_j}{(\boldsymbol{\beta}'\boldsymbol{\beta})^2}\sum_i a_i + 2\frac{\sqrt{a_j}}{\boldsymbol{\beta}'\boldsymbol{\beta}}\sum_i \sqrt{a_i}(\hat{\boldsymbol{\epsilon}}_i'\hat{\boldsymbol{\epsilon}}_j + \mathrm{Cov}(\mathbf{E})_{ij}) + \sum_i (\hat{\boldsymbol{\epsilon}}_i'\hat{\boldsymbol{\epsilon}}_j + \mathrm{Cov}(\mathbf{E})_{ij})^2. \qquad (4)
\end{aligned}$$

The middle line above uses the orthogonality (by definition) of $\boldsymbol{\beta}$ and $\epsilon_j$. As Eq. 1 in the main text does not accommodate dependence between $a_j$ and $\sum_i r_{ij}^2$, resulting estimates of $h^2_{\mathrm{SNP}}$ are in general biased. As $\sqrt{a_i}$, and $\hat{\boldsymbol{\epsilon}}_i'\hat{\boldsymbol{\epsilon}}_j + \mathrm{Cov}(\mathbf{E})_{ij}$ can be either positive or negative, the sum associated with $\sqrt{a_j}$ will tend to cancel so that LD score is almost always positively correlated with $a_j$, which in LDSC will inflate estimates of $h^2_{\mathrm{SNP}}$. As $\sum_i a_i = \sum_i \mathrm{Cov}(\mathbf{Z}_i, \mathbf{X}\boldsymbol{\beta})^2$, the magnitude of inflation increases with the square correlations between the genotypes $\mathbf{Z}$ and the ignored confounding effect $\mathbf{X}\boldsymbol{\beta}$ as well as with the proportion of phenotypic variation attributed to the confounding effect.

To aid intuition about this effect, consider the widespread and successful use of principal components of SNP genotypes to adjust for confounding due to population structure. In order to maximise variation explained, the first principal component usually has high loadings from sets of SNPs in high LD and it usually captures a large component of confounding, supporting our claim of an association between LD and confounding. If the stratification is due to ignored sub-populations, SNPs with greater allele frequency differences between sub-populations will have greater induced LD, so that higher LD score will be associated with the SNPs more predictive of sub-population membership.

To illustrate the impact of the LD/confounding relationship on estimation of $h^2_{\mathrm{SNP}}$, we extracted the confounding effects $\mathbf{X}\boldsymbol{\beta}$ from our simulations, and treated their standardised values as the phenotype in a GWAS analysis. The resulting estimate of $h^2_{\mathrm{SNP}}$ was multiplied by 0.1 to have the same scale as the confounding component in a $C1$ phenotype. The positive relationship between LD and confounding generates positive estimates of $h^2_{\mathrm{SNP}}$ (Figure S10), although these estimates tend to be lower than the observed inflation in $h^2_{\mathrm{SNP}}$ estimates in the original analyses. This is consistent with the expected cross-product of the confounding effect test statistic and confounding-free phenotype test statistic also showing a positive relationship with LD score.

### 3 Approximate expectations of association statistics

In this section, we will provide the working for determining expected association statistics for a wide range of possible GWAS design analysed with fixed effect linear regression. These scenarios cover when $\mathbf{X}$ was and was not included in the regression, and whether $\mathbf{X}$ is associated with $\mathbf{Z}$.

#### 3.1 Single-SNP analysis with no covariates fitted and Z known

In single-SNP regression of a quantitative phenotype,

$$T_j^2 = \frac{(\mathbf{y}-\bar{\mathbf{y}}\mathbf{1})'\mathbf{Z}_j(\mathbf{Z}_j'\mathbf{Z}_j)^{-1}\mathbf{Z}_j'(\mathbf{y}-\bar{\mathbf{y}}\mathbf{1})}{(\mathbf{y}-\bar{\mathbf{y}}\mathbf{1})'(\mathbf{I}-\mathbf{Z}_j(\mathbf{Z}_j'\mathbf{Z}_j)^{-1}\mathbf{Z}_j')(\mathbf{y}-\bar{\mathbf{y}}\mathbf{1})/(n-2)}. \tag{5}$$

Following Bulik-Sullivan et al. (2015a), we consider the numerator and denominator of (5) separately. Some notation: $\mathbf{J} = \mathbf{11}'$ is a matrix of ones, so that $\mathbf{J}^2 = n\mathbf{J}$, and Tr denotes the trace of a matrix. In particular if $\mathbf{a}$ is a vector, $\mathrm{Tr}(\mathbf{aa}') = \mathbf{a}'\mathbf{a}$. We require

$$\begin{aligned}
\mathrm{E}[(\mathbf{y}-\bar{\mathbf{y}}\mathbf{1}))(\mathbf{y}-\bar{\mathbf{y}}\mathbf{1})'] &= (\mathbf{I}-\mathbf{J}/n)\mathrm{E}[\mathbf{yy}'](\mathbf{I}-\mathbf{J}/n)' \\
&= (\mathbf{I}-\mathbf{J}/n)(\mathbf{Z\Sigma Z}' + \sigma_e^2\mathbf{I} + \mathbf{X}\boldsymbol{\beta}\boldsymbol{\beta}'\mathbf{X}')(\mathbf{I}-\mathbf{J}/n)' \\
&= \mathbf{Z\Sigma Z}' + \sigma_e^2(\mathbf{I} - 2\mathbf{J}/n + \mathbf{JJ}/n^2) + \mathbf{X}\boldsymbol{\beta}\boldsymbol{\beta}'\mathbf{X}' \qquad (6) \\
&= \mathbf{Z\Sigma Z}' + \sigma_e^2(\mathbf{I} - \mathbf{J}/n) + \mathbf{X}\boldsymbol{\beta}\boldsymbol{\beta}'\mathbf{X}' \qquad (7)
\end{aligned}$$

with the final two equalities following from the standardisation of $\mathbf{Z}$, which implies $\mathbf{Z}'\mathbf{J} = \mathbf{0}_{m\times n}$ and standardisation of $\mathbf{X}$ implying $\mathbf{X}'\mathbf{J} = \mathbf{0}_{p\times n}$. Using (7), the expected value of the numerator of (5), which following ANOVA conventions we call $SSR_j$, is

$$\begin{aligned}
\mathrm{E}[SSR_j|\mathbf{Z},\mathbf{X}] &= \mathrm{E}[(\mathbf{y}-\bar{\mathbf{y}}\mathbf{1})'\mathbf{Z}_j(\mathbf{Z}_j'\mathbf{Z}_j)^{-1}\mathbf{Z}_j'(\mathbf{y}-\bar{\mathbf{y}}\mathbf{1})] \\
&= \mathrm{Tr}(\mathbf{Z}_j(\mathbf{Z}_j'\mathbf{Z}_j)^{-1}\mathbf{Z}_j'\mathrm{E}[(\mathbf{y}-\bar{\mathbf{y}}\mathbf{1}))(\mathbf{y}-\bar{\mathbf{y}}\mathbf{1})']) \\
&= \mathrm{Tr}(\mathbf{Z}_j(\mathbf{Z}_j'\mathbf{Z}_j)^{-1}\mathbf{Z}_j'\{\mathbf{Z\Sigma Z}' + \sigma_e^2(\mathbf{I}_n - \mathbf{J}/n) + \mathbf{X}\boldsymbol{\beta}\boldsymbol{\beta}'\mathbf{X}'\}) \\
&= \mathrm{Tr}(\mathbf{Z}_j(\mathbf{Z}_j'\mathbf{Z}_j)^{-1}\mathbf{Z}_j'\{\mathbf{Z\Sigma Z}' + \sigma_e^2\mathbf{I}_n + \mathbf{X}\boldsymbol{\beta}\boldsymbol{\beta}'\mathbf{X}'\}) \\
&= \mathrm{Tr}((\mathbf{Z}_j'\mathbf{Z}_j)^{-1}\mathbf{Z}_j'\mathbf{Z\Sigma Z}'\mathbf{Z}_j + \sigma_e^2(\mathbf{Z}_j'\mathbf{Z}_j)^{-1}(\mathbf{Z}_j'\mathbf{Z}_j) + (\mathbf{Z}_j'\mathbf{Z}_j)^{-1}\mathbf{Z}_j'\mathbf{X}\boldsymbol{\beta}\boldsymbol{\beta}'\mathbf{X}'\mathbf{Z}_j\}) \\
&= (\mathbf{Z}_j'\mathbf{Z\Sigma Z}'\mathbf{Z}_j + \mathbf{Z}_j'\mathbf{X}\boldsymbol{\beta}\boldsymbol{\beta}'\mathbf{X}'\mathbf{Z}_j)/(n-1) + \sigma_e^2. \qquad (8)
\end{aligned}$$

Note that $\mathbf{Z}_j'\mathbf{Z} = (n-1)\hat{\mathrm{Cor}}(\mathbf{Z}_j, \mathbf{Z})$, and if $\mathbf{X}$ is standardised, then $\mathbf{Z}_j'\mathbf{X} = (n-1)\hat{\mathrm{Cor}}(\mathbf{Z}_j, \mathbf{X})$. Since $\mathbf{\Sigma}$ is diagonal, we can write

$$\begin{aligned}
\mathrm{E}[SSR_j|\mathbf{Z},\mathbf{X}] &= (n-1)\left\{\sum_{i=1}^{m}\hat{r}_{ij}^2\sigma_i^2+\left(\sum_{k=1}^{p}\hat{\mathrm{Cor}}(\mathbf{Z}_j,\mathbf{X}_k)\boldsymbol{\beta}_k\right)^2\right\}+\sigma_e^2 \\
&= \sigma_e^2+(n-1)\sigma_j^2+(n-1)\sum_{i\neq j}\hat{r}_{ij}^2\sigma_i^2+(n-1)\left(\sum_{k=1}^{p}\hat{\mathrm{Cor}}(\mathbf{Z}_j,\mathbf{X}_k)\boldsymbol{\beta}_k\right)^2 \qquad (9)
\end{aligned}$$

where $\hat{r}_{ij}=\hat{\mathrm{Cor}}(\mathbf{Z}_i,\mathbf{Z}_j)$. Write $SSE_j$ for $n-2$ times the denominator of (5). Then

$$\begin{aligned}
\mathrm{E}[SSE_j|\mathbf{Z},\mathbf{X}] &= \mathrm{E}[(\mathbf{y}-\bar{\mathbf{y}}\mathbf{1})'(\mathbf{I}-\mathbf{Z}_j(\mathbf{Z}_j'\mathbf{Z}_j)^{-1}\mathbf{Z}_j')(\mathbf{y}-\bar{\mathbf{y}}\mathbf{1})] \\
&= \mathrm{E}[(\mathbf{y}-\bar{\mathbf{y}}\mathbf{1})'(\mathbf{y}-\bar{\mathbf{y}}\mathbf{1})]-\mathrm{E}[SSR|\mathbf{Z},\mathbf{X}] \\
&= \mathrm{Tr}(\mathrm{E}[(\mathbf{y}-\bar{\mathbf{y}}\mathbf{1})\mathbf{y}-\bar{\mathbf{y}}\mathbf{1})'])-\mathrm{E}[SSR|\mathbf{Z},\mathbf{X}] \\
&= \mathrm{Tr}(\mathbf{Z}\boldsymbol{\Sigma}\mathbf{Z}'+\sigma_e^2(\mathbf{I}-\mathbf{J}/n)+\mathbf{X}\boldsymbol{\beta}\boldsymbol{\beta}'\mathbf{X}')-\mathrm{E}[SSR_j|\mathbf{Z},\mathbf{X}] \\
&= \mathrm{Tr}(\mathbf{Z}'\mathbf{Z}\boldsymbol{\Sigma})+(n-1)\sigma_e^2+\mathrm{Tr}(\boldsymbol{\beta}'\mathbf{X}'\mathbf{X}\boldsymbol{\beta})-\mathrm{E}[SSR_j|\mathbf{Z},\mathbf{X}] \\
&= (n-1)\left\{\sum_{i=1}^{m}\sigma_j^2+\sigma_e^2+\boldsymbol{\beta}'\hat{\mathrm{Var}}[\mathbf{X}]\boldsymbol{\beta}\right\}-\sigma_e^2-(n-1)\left\{\sum_{i=1}^{m}\hat{r}_{ij}^2\sigma_i^2-\left(\sum_{k=1}^{p}\hat{\mathrm{Cor}}(\mathbf{Z}_j,\mathbf{X}_k)\boldsymbol{\beta}_k\right)^2\right\} \\
&= (n-2)\sigma_e^2+(n-1)\left\{\sum_{i\neq j}(1-\hat{r}_{ij}^2)\sigma_i^2+\sum_{k=1}^{p}\sum_{l=1}^{p}(\hat{\mathrm{Var}}[\mathbf{X}]_{kl}-\hat{\mathrm{Cor}}(\mathbf{Z}_j,\mathbf{X}_k)\hat{\mathrm{Cor}}(\mathbf{Z}_j,\mathbf{X}_l))\boldsymbol{\beta}_k\boldsymbol{\beta}_l\right\} \qquad (10)
\end{aligned}$$

#### 3.2 Single-SNP analysis with no covariates fitted and Z unknown

Usually, GWAS results will not include the SNP sample correlation matrix, which is required for the expectations in section 3.1. Rather, it is estimated from a reference panel. Rather than use $\hat{r}_{ij}^2$ directly, Bulik-Sullivan et al. (2015a) used a transformation to reduce bias: $r_{ij}^2=((n-1)\hat{r}_{ij}^2-1)/(n-2)$. Substituting this into (9) for SNP-SNP and SNP-covariate correlations, we obtain

$$\begin{aligned}
\mathrm{E}[SSR_j|\mathbf{X}] &= \sigma_e^2+(n-1)\sigma_j^2+(n-1)\sum_{i\neq j}\left\{\frac{(n-2)r_{ij}^2}{n-1}+\frac{1}{n-1}\right\}\sigma_j^2+(n-1)\left(\sum_{k=1}^{p}\hat{\mathrm{Cor}}(\mathbf{Z}_j,\mathbf{X}_k)\boldsymbol{\beta}_k\right)^2 \\
&= \sigma_e^2+\sum_{i=1}^{m}\sigma_j^2+(n-2)\sigma_j^2+(n-2)\sum_{i\neq j}r_{ij}^2\sigma_i^2+(n-1)\sigma_c^2\hat{\mathrm{Cor}}(\mathbf{Z}_j,\mathbf{X}\boldsymbol{\beta})^2 \\
&= \sigma_e^2+\sum_{i=1}^{m}\sigma_j^2+(n-2)\sigma_j^2+(n-2)\sum_{i\neq j}r_{ij}^2\sigma_i^2+(n-1)\sigma_c^2\left\{\frac{(n-2)\mathrm{Cor}(\mathbf{Z}_j,\mathbf{X}\boldsymbol{\beta})^2}{n-1}+\frac{1}{n-1}\right\} \\
&= \sigma_y^2+\sigma_c^2+(n-2)\sigma_j^2+(n-2)\sum_{i\neq j}r_{ij}^2\sigma_i^2+(n-2)\left(\sum_{k=1}^{p}\mathrm{Cor}(\mathbf{Z}_j,\mathbf{X}_k)\boldsymbol{\beta}_k\right)^2. \qquad (11)
\end{aligned}$$

For large $n$,

$$
\begin{aligned}
\mathrm{E}[SSR_j|\mathbf{X}] &\approx \sigma_y^2+\sigma_c^2+n\sigma_j^2+n\sum_{i\neq j} r_{ij}^2\sigma_i^2+n\sum_{k=1}^{p}\sum_{l=1}^{p}\mathrm{Cor}(\mathbf{Z}_j,\mathbf{X}_k)\mathrm{Cor}(\mathbf{Z}_j,\mathbf{X}_l)\boldsymbol{\beta}_k\boldsymbol{\beta}_l. \qquad (12)\\
\mathrm{E}[SSE_j|\mathbf{X}] &= \mathrm{E}[\mathrm{E}[SSE_j|\mathbf{Z},\mathbf{X}]]=\mathrm{E}[\mathrm{E}[SST|\mathbf{Z},\mathbf{X}]]-\mathrm{E}[\mathrm{E}[SSR_j|\mathbf{Z},\mathbf{X}]]\\
&= (n-2)\left[\sigma_y^2+\sigma_c^2-\sigma_j^2-\sum_{i\neq j} r_{ij}^2\sigma_{\boldsymbol{\alpha}_i}^2-\left(\sum_{k=1}^{p}\mathrm{Cor}(\mathbf{Z}_j,\mathbf{X}_k)\boldsymbol{\beta}_k\right)^2\right] \qquad (13)\\
&= (n-2)c_j(\sigma_y^2+\sigma_c^2), \qquad (14)
\end{aligned}
$$

where $c_j=1-\sum_i r_{ij}^2h_{i,\mathrm{b}}^2-(\sum_{k=1}^{p}\mathrm{Cor}(\mathbf{Z}_j,\mathbf{X}_k)\boldsymbol{\beta}_k)^2/(\sigma_y^2+\sigma_c^2)$.

#### 3.3 Single-SNP analysis with covariates fitted and Z known

The sum of squares required to determine expectations when covariates are fitted corresponds to quantities required for a partial $F$ test comparing the fit of the nested models:

$$
\begin{aligned}
\mathbf{y} &= \mathbf{1}\mu+\mathbf{X}\boldsymbol{\beta}+\boldsymbol{\varepsilon} \qquad (15)\\
\mathbf{y} &= \mathbf{1}\mu+\mathbf{X}\boldsymbol{\beta}+\mathbf{Z}_j\boldsymbol{\gamma}_j+\boldsymbol{\varepsilon}, \qquad (16)
\end{aligned}
$$

where $\boldsymbol{\beta}$, $\boldsymbol{\gamma}_j$ are fixed and $\varepsilon\sim\mathcal{N}(\mathbf{0},\sigma_e^2\mathbf{I})$. We use subscripts 1 and 2 to denote models (15) and (16).

$$
\begin{aligned}
\mathrm{E}[SSR_{j,1}|\mathbf{Z},\mathbf{X}] &= \mathrm{E}[(\mathbf{y}-\bar{\mathbf{y}}\mathbf{1})'\mathbf{X}(\mathbf{X}'\mathbf{X})^{-1}\mathbf{X}'(\mathbf{y}-\bar{\mathbf{y}}\mathbf{1})]\\
&= \mathrm{Tr}(\mathbf{X}(\mathbf{X}'\mathbf{X})^{-1}\mathbf{X}'\mathrm{E}[(\mathbf{y}-\bar{\mathbf{y}}\mathbf{1}))(\mathbf{y}-\bar{\mathbf{y}}\mathbf{1})'])\\
&= \mathrm{Tr}(\mathbf{X}(\mathbf{X}'\mathbf{X})^{-1}\mathbf{X}'\{\mathbf{Z}\boldsymbol{\Sigma}\mathbf{Z}'+\sigma_e^2\mathbf{I}_n-\frac{\sigma_e^2}{n}\mathbf{J}+\mathbf{X}\boldsymbol{\beta}\boldsymbol{\beta}'\mathbf{X}'\})\\
&= \mathrm{Tr}(\mathbf{X}(\mathbf{X}'\mathbf{X})^{-1}\mathbf{X}'\{\mathbf{Z}\boldsymbol{\Sigma}\mathbf{Z}'+\sigma_e^2\mathbf{I}_n+\mathbf{X}\boldsymbol{\beta}\boldsymbol{\beta}'\mathbf{X}'\})\\
&= \mathrm{Tr}((\mathbf{X}'\mathbf{X})^{-1}\mathbf{X}'\mathbf{Z}\boldsymbol{\Sigma}\mathbf{Z}'\mathbf{X}+\sigma_e^2(\mathbf{X}'\mathbf{X})^{-1}\mathbf{X}'\mathbf{X}+(\mathbf{X}'\mathbf{X})^{-1}\mathbf{X}'\mathbf{X}\boldsymbol{\beta}\boldsymbol{\beta}'\mathbf{X}'\mathbf{X}\})\\
&= \mathrm{Tr}((\mathbf{X}'\mathbf{X})^{-1}\mathbf{X}'\mathbf{Z}\boldsymbol{\Sigma}\mathbf{Z}'\mathbf{X})+p\sigma_e^2+\boldsymbol{\beta}'\mathbf{X}'\mathbf{X}\boldsymbol{\beta}. \qquad (17)
\end{aligned}
$$

$$
\begin{aligned}
\mathrm{E}[SSR_{j,2}|\mathbf{Z},\mathbf{X}] &= \mathrm{E}\left[(\mathbf{y}-\bar{\mathbf{y}}\mathbf{1})'\begin{pmatrix}\mathbf{X} & \mathbf{Z}_j\end{pmatrix}\begin{pmatrix}\mathbf{X}'\mathbf{X} & \mathbf{X}'\mathbf{Z}_j\\ \mathbf{Z}_j'\mathbf{X} & \mathbf{Z}_j'\mathbf{Z}_j\end{pmatrix}^{-1}\begin{pmatrix}\mathbf{X}'\\ \mathbf{Z}_j'\end{pmatrix}(\mathbf{y}-\bar{\mathbf{y}}\mathbf{1})\right]\\
&= \mathrm{Tr}\left(\begin{pmatrix}\mathbf{X}'\mathbf{X} & \mathbf{X}'\mathbf{Z}_j\\ \mathbf{Z}_j'\mathbf{X} & \mathbf{Z}_j'\mathbf{Z}_j\end{pmatrix}^{-1}\begin{pmatrix}\mathbf{X}'\\ \mathbf{Z}_j'\end{pmatrix}\mathrm{E}[(\mathbf{y}-\bar{\mathbf{y}}\mathbf{1}))(\mathbf{y}-\bar{\mathbf{y}}\mathbf{1})']\begin{pmatrix}\mathbf{X} & \mathbf{Z}_j\end{pmatrix}\right)
\end{aligned}
$$

$$\begin{aligned} &= \operatorname{Tr}\left( \begin{pmatrix} \mathbf{X}'\mathbf{X} & \mathbf{X}'\mathbf{Z}_j \\ \mathbf{Z}_j'\mathbf{X} & \mathbf{Z}_j'\mathbf{Z}_j \end{pmatrix}^{-1} \begin{pmatrix} \mathbf{X}' \\ \mathbf{Z}_j' \end{pmatrix} \{\mathbf{Z}\boldsymbol{\Sigma}\mathbf{Z}' + \sigma_e^2\mathbf{I}_n + \mathbf{X}\boldsymbol{\beta}\boldsymbol{\beta}'\mathbf{X}'\} \begin{pmatrix} \mathbf{X} & \mathbf{Z}_j \end{pmatrix} \right) \\ &= \operatorname{Tr}\left( \begin{pmatrix} \mathbf{X}'\mathbf{X} & \mathbf{X}'\mathbf{Z}_j \\ \mathbf{Z}_j'\mathbf{X} & \mathbf{Z}_j'\mathbf{Z}_j \end{pmatrix}^{-1} \begin{pmatrix} \mathbf{X}'\mathbf{Z}\boldsymbol{\Sigma}\mathbf{Z}'\mathbf{X} & \mathbf{X}'\mathbf{Z}\boldsymbol{\Sigma}\mathbf{Z}'\mathbf{Z}_j \\ \mathbf{Z}_j'\mathbf{Z}\boldsymbol{\Sigma}\mathbf{Z}'\mathbf{X} & \mathbf{Z}_j'\mathbf{Z}\boldsymbol{\Sigma}\mathbf{Z}'\mathbf{Z}_j \end{pmatrix} \right) + (p+1)\sigma_e^2 \\ &\quad + \operatorname{Tr}\left( \begin{pmatrix} \mathbf{X}'\mathbf{X} & \mathbf{X}'\mathbf{Z}_j \\ \mathbf{Z}_j'\mathbf{X} & \mathbf{Z}_j'\mathbf{Z}_j \end{pmatrix}^{-1} \begin{pmatrix} \mathbf{X}'\mathbf{X}\boldsymbol{\beta}\boldsymbol{\beta}'\mathbf{X}'\mathbf{X} & \mathbf{X}'\mathbf{X}\boldsymbol{\beta}\boldsymbol{\beta}'\mathbf{X}'\mathbf{Z}_j \\ \mathbf{Z}_j'\mathbf{X}\boldsymbol{\beta}\boldsymbol{\beta}'\mathbf{X}'\mathbf{X} & \mathbf{Z}_j'\mathbf{X}\boldsymbol{\beta}\boldsymbol{\beta}'\mathbf{X}'\mathbf{Z}_j \end{pmatrix} \right). \end{aligned} \tag{18}$$

To simplify, we expand the inverse in (18) using the block inversion formula, obtaining

$$\begin{pmatrix} \mathbf{X}'\mathbf{X} & \mathbf{X}'\mathbf{Z}_j \\ \mathbf{Z}_j'\mathbf{X} & \mathbf{Z}_j'\mathbf{Z}_j \end{pmatrix}^{-1} = \begin{pmatrix} (\mathbf{X}'\mathbf{X})^{-1} + (\mathbf{X}'\mathbf{X})^{-1}\mathbf{X}'\mathbf{Z}_j\mathbf{B}_j^{-1}\mathbf{Z}_j'\mathbf{X}(\mathbf{X}'\mathbf{X})^{-1} & -(\mathbf{X}'\mathbf{X})^{-1}\mathbf{X}'\mathbf{Z}_j\mathbf{B}_j^{-1} \\ -\mathbf{B}_j^{-1}\mathbf{Z}_j'\mathbf{X}(\mathbf{X}'\mathbf{X})^{-1} & \mathbf{B}_j^{-1} \end{pmatrix}, \tag{19}$$

where $\mathbf{B}_j = \mathbf{Z}_j'(\mathbf{I} - \mathbf{X}(\mathbf{X}'\mathbf{X})^{-1}\mathbf{X}')\mathbf{Z}_j$. Substituting (19) into (18) and noting that only the diagonal blocks of (18) are of interest, we find for the blocks associated with $\mathbf{Z}\boldsymbol{\Sigma}\mathbf{Z}'$,

$$\begin{aligned} &\begin{pmatrix} (\mathbf{X}'\mathbf{X})^{-1} + (\mathbf{X}'\mathbf{X})^{-1}\mathbf{X}'\mathbf{Z}_j\mathbf{B}_j^{-1}\mathbf{Z}_j'\mathbf{X}(\mathbf{X}'\mathbf{X})^{-1} & -(\mathbf{X}'\mathbf{X})^{-1}\mathbf{X}'\mathbf{Z}_j\mathbf{B}_j^{-1} \end{pmatrix} \begin{pmatrix} \mathbf{X}'\mathbf{Z}\boldsymbol{\Sigma}\mathbf{Z}'\mathbf{X} \\ \mathbf{Z}_j'\mathbf{Z}\boldsymbol{\Sigma}\mathbf{Z}'\mathbf{X} \end{pmatrix} \\ &\qquad = (\mathbf{X}'\mathbf{X})^{-1}\mathbf{X}'\mathbf{Z}\boldsymbol{\Sigma}\mathbf{Z}'\mathbf{X} - (\mathbf{X}'\mathbf{X})^{-1}\mathbf{X}'\mathbf{Z}_j\mathbf{B}_j^{-1}\mathbf{Z}_j'(\mathbf{I} - \mathbf{X}(\mathbf{X}'\mathbf{X})^{-1}\mathbf{X}')\mathbf{Z}\boldsymbol{\Sigma}\mathbf{Z}'\mathbf{X} \end{aligned} \tag{20}$$

$$\begin{pmatrix} -\mathbf{B}_j^{-1}\mathbf{Z}_j'\mathbf{X}(\mathbf{X}'\mathbf{X})^{-1} & \mathbf{B}_j^{-1} \end{pmatrix} \begin{pmatrix} \mathbf{X}'\mathbf{Z}\boldsymbol{\Sigma}\mathbf{Z}'\mathbf{Z}_j \\ \mathbf{Z}_j'\mathbf{Z}\boldsymbol{\Sigma}\mathbf{Z}'\mathbf{Z}_j \end{pmatrix} = \mathbf{B}_j^{-1}\mathbf{Z}_j'(\mathbf{I} - \mathbf{X}(\mathbf{X}'\mathbf{X})^{-1}\mathbf{X}')\mathbf{Z}\boldsymbol{\Sigma}\mathbf{Z}'\mathbf{Z}_j. \tag{21}$$

For the blocks associated with $\mathbf{X}\boldsymbol{\beta}\boldsymbol{\beta}'\mathbf{X}'$, we get

$$\begin{aligned} &\begin{pmatrix} (\mathbf{X}'\mathbf{X})^{-1} + (\mathbf{X}'\mathbf{X})^{-1}\mathbf{X}'\mathbf{Z}_j\mathbf{B}_j^{-1}\mathbf{Z}_j'\mathbf{X}(\mathbf{X}'\mathbf{X})^{-1} & -(\mathbf{X}'\mathbf{X})^{-1}\mathbf{X}'\mathbf{Z}_j\mathbf{B}_j^{-1} \end{pmatrix} \begin{pmatrix} \mathbf{X}'\mathbf{X}\boldsymbol{\beta}\boldsymbol{\beta}'\mathbf{X}'\mathbf{X} \\ \mathbf{Z}_j'\mathbf{X}\boldsymbol{\beta}\boldsymbol{\beta}'\mathbf{X}'\mathbf{X} \end{pmatrix} \\ &\qquad = \boldsymbol{\beta}\boldsymbol{\beta}'\mathbf{X}'\mathbf{X} + (\mathbf{X}'\mathbf{X})^{-1}\mathbf{X}'\mathbf{Z}_j\mathbf{B}_j^{-1}\mathbf{Z}_j'\mathbf{X}\boldsymbol{\beta}\boldsymbol{\beta}'\mathbf{X}'\mathbf{X} - (\mathbf{X}'\mathbf{X})^{-1}\mathbf{X}'\mathbf{Z}_j\mathbf{B}_j^{-1}\mathbf{Z}_j'\mathbf{X}\boldsymbol{\beta}\boldsymbol{\beta}'\mathbf{X}'\mathbf{X} = \boldsymbol{\beta}\boldsymbol{\beta}'\mathbf{X}'\mathbf{X} \end{aligned} \tag{22}$$

$$\begin{pmatrix} -\mathbf{B}_j^{-1}\mathbf{Z}_j'\mathbf{X}(\mathbf{X}'\mathbf{X})^{-1} & \mathbf{B}_j^{-1} \end{pmatrix} \begin{pmatrix} \mathbf{X}'\mathbf{X}\boldsymbol{\beta}\boldsymbol{\beta}'\mathbf{X}'\mathbf{Z}_j \\ \mathbf{Z}_j'\mathbf{X}\boldsymbol{\beta}\boldsymbol{\beta}'\mathbf{X}'\mathbf{Z}_j \end{pmatrix} = -\mathbf{B}_j^{-1}\mathbf{Z}_j'\mathbf{X}\boldsymbol{\beta}\boldsymbol{\beta}'\mathbf{X}'\mathbf{Z}_j + \mathbf{B}_j^{-1}\mathbf{Z}_j'\mathbf{X}\boldsymbol{\beta}\boldsymbol{\beta}'\mathbf{X}'\mathbf{Z}_j = 0 \tag{23}$$

Substituting (20)-(23) into (18), we obtain

$$\begin{aligned} \mathrm{E}[SSR_{j,2}|\mathbf{Z},\mathbf{X}] &= \operatorname{Tr}((\mathbf{X}'\mathbf{X})^{-1}\mathbf{X}'\mathbf{Z}\boldsymbol{\Sigma}\mathbf{Z}'\mathbf{X} - (\mathbf{X}'\mathbf{X})^{-1}\mathbf{X}'\mathbf{Z}_j\mathbf{B}_j^{-1}\mathbf{Z}_j'(\mathbf{I} - \mathbf{X}(\mathbf{X}'\mathbf{X})^{-1}\mathbf{X}')\mathbf{Z}\boldsymbol{\Sigma}\mathbf{Z}'\mathbf{X}) \\ &\quad + \operatorname{Tr}(\mathbf{B}_j^{-1}\mathbf{Z}_j'(\mathbf{I} - \mathbf{X}(\mathbf{X}'\mathbf{X})^{-1}\mathbf{X}')\mathbf{Z}\boldsymbol{\Sigma}\mathbf{Z}'\mathbf{Z}_j) + (p+1)\sigma_e^2 + \operatorname{Tr}(\boldsymbol{\beta}\boldsymbol{\beta}'\mathbf{X}'\mathbf{X}) \\ &= \operatorname{Tr}((\mathbf{X}'\mathbf{X})^{-1}\mathbf{X}'\mathbf{Z}\boldsymbol{\Sigma}\mathbf{Z}'\mathbf{X}) - \operatorname{Tr}(\mathbf{X}(\mathbf{X}'\mathbf{X})^{-1}\mathbf{X}'\mathbf{Z}_j\mathbf{B}_j^{-1}\mathbf{Z}_j'(\mathbf{I} - \mathbf{X}(\mathbf{X}'\mathbf{X})^{-1}\mathbf{X}')\mathbf{Z}\boldsymbol{\Sigma}\mathbf{Z}') \\ &\quad + \operatorname{Tr}(\mathbf{Z}_j\mathbf{B}_j^{-1}\mathbf{Z}_j'(\mathbf{I} - \mathbf{X}(\mathbf{X}'\mathbf{X})^{-1}\mathbf{X}')\mathbf{Z}\boldsymbol{\Sigma}\mathbf{Z}') + (p+1)\sigma_e^2 + \boldsymbol{\beta}'\mathbf{X}'\mathbf{X}\boldsymbol{\beta} \end{aligned}$$

$$= \text{Tr}((\mathbf{X}'\mathbf{X})^{-1}\mathbf{X}'\mathbf{Z}\boldsymbol{\Sigma}\mathbf{Z}'\mathbf{X}) + \text{Tr}((\mathbf{I}-\mathbf{X}(\mathbf{X}'\mathbf{X})^{-1}\mathbf{X}')\mathbf{Z}_j\mathbf{B}_j^{-1}\mathbf{Z}_j'(\mathbf{I}-\mathbf{X}(\mathbf{X}'\mathbf{X})^{-1}\mathbf{X}')\mathbf{Z}\boldsymbol{\Sigma}\mathbf{Z}') + (p+1)\sigma_e^2 + \boldsymbol{\beta}'\mathbf{X}'\mathbf{X}\boldsymbol{\beta}. \quad (24)$$

The difference in regression sum of squares between model 1 (15) and model 2 (16) is

$$\begin{aligned} \text{E}[SSR_{j,2} - SSR_{j,1}|\mathbf{Z},\mathbf{X}] &= \text{Tr}((\mathbf{I}-\mathbf{X}(\mathbf{X}'\mathbf{X})^{-1}\mathbf{X}')\mathbf{Z}_j\mathbf{B}_j^{-1}\mathbf{Z}_j'(\mathbf{I}-\mathbf{X}(\mathbf{X}'\mathbf{X})^{-1}\mathbf{X}')\mathbf{Z}\boldsymbol{\Sigma}\mathbf{Z}') + (p+1)\sigma_e^2 - p\sigma_e^2 \\ &\quad + \text{Tr}((\mathbf{X}'\mathbf{X})^{-1}\mathbf{X}'\mathbf{Z}\boldsymbol{\Sigma}\mathbf{Z}'\mathbf{X}) - \text{Tr}((\mathbf{X}'\mathbf{X})^{-1}\mathbf{X}'\mathbf{Z}\boldsymbol{\Sigma}\mathbf{Z}'\mathbf{X}) + \boldsymbol{\beta}'\mathbf{X}'\mathbf{X}\boldsymbol{\beta} - \boldsymbol{\beta}'\mathbf{X}'\mathbf{X}\boldsymbol{\beta} \\ &= \sigma_e^2 + \text{Tr}((\mathbf{I}-\mathbf{X}(\mathbf{X}'\mathbf{X})^{-1}\mathbf{X}')\mathbf{Z}_j\mathbf{B}_j^{-1}\mathbf{Z}_j'(\mathbf{I}-\mathbf{X}(\mathbf{X}'\mathbf{X})^{-1}\mathbf{X}')\mathbf{Z}\boldsymbol{\Sigma}\mathbf{Z}'), \qquad (25) \end{aligned}$$

and the error sum of squares for model 2 is

$$\begin{aligned} \text{E}[SSE_{j,2}|\mathbf{Z},\mathbf{X}] &= \text{E}[(\mathbf{y}-\bar{\mathbf{y}}\mathbf{1})'(\mathbf{y}-\bar{\mathbf{y}}\mathbf{1})] - \text{E}[SSR_{j,2}|\mathbf{Z},\mathbf{X}] \\ &= \text{Tr}(\mathbf{Z}\boldsymbol{\Sigma}\mathbf{Z}') + (n-1)\sigma_e^2 + \text{Tr}(\mathbf{X}\boldsymbol{\beta}\boldsymbol{\beta}'\mathbf{X}') - \text{E}[SSR_2|\mathbf{Z},\mathbf{X}] \\ &= \text{Tr}(\mathbf{Z}\boldsymbol{\Sigma}\mathbf{Z}') + (n-1)\sigma_e^2 + \boldsymbol{\beta}'\mathbf{X}'\mathbf{X}\boldsymbol{\beta} - \text{Tr}(\mathbf{X}(\mathbf{X}'\mathbf{X})^{-1}\mathbf{X}'\mathbf{Z}\boldsymbol{\Sigma}\mathbf{Z}') \\ &\quad - \text{Tr}((\mathbf{I}-\mathbf{X}(\mathbf{X}'\mathbf{X})^{-1}\mathbf{X}')\mathbf{Z}_j\mathbf{B}_j^{-1}\mathbf{Z}_j'(\mathbf{I}-\mathbf{X}(\mathbf{X}'\mathbf{X})^{-1}\mathbf{X}')\mathbf{Z}\boldsymbol{\Sigma}\mathbf{Z}') - (p+1)\sigma_e^2 - \boldsymbol{\beta}'\mathbf{X}'\mathbf{X}\boldsymbol{\beta} \\ &= \text{Tr}((\mathbf{I}-\mathbf{X}(\mathbf{X}'\mathbf{X})^{-1}\mathbf{X}')\mathbf{Z}\boldsymbol{\Sigma}\mathbf{Z}') - \text{Tr}((\mathbf{I}-\mathbf{X}(\mathbf{X}'\mathbf{X})^{-1}\mathbf{X}')\mathbf{Z}_j\mathbf{B}_j^{-1}\mathbf{Z}_j'(\mathbf{I}-\mathbf{X}(\mathbf{X}'\mathbf{X})^{-1}\mathbf{X}')\mathbf{Z}\boldsymbol{\Sigma}\mathbf{Z}') \\ &= (n-p-2)\sigma_e^2 \qquad (26) \end{aligned}$$

#### 3.4 Expected sums of squares when $\mathbf{Z}, \mathbf{X}$ unknown and independent

Having marginalised expectations over $\mathbf{Z}$ in Section 3.2, the next step is to marginalise over $\mathbf{X}$ in order to deal with the case when $\mathbf{X}$ is not available. Since there are usually no reference panels of covariates, stronger assumptions are needed than for $\mathbf{Z}$. One plausible assumption is that $\mathbf{X}$ and $\mathbf{Z}$ are independent, so the $\mathbf{X}$ includes covariates and not confounders. In addition, and without loss of generality, we will assume that the columns of $\mathbf{X}$ are mutually independent.

##### 3.4.1 Single-SNP analysis with no covariates fitted, $\text{Cor}(\mathbf{X}, \mathbf{Z}) = 0$.

The expected sum of squares when $\mathbf{X}$ is independent of $\mathbf{Z}$ and was included in the analysis model correspond to (12) and (14) with $\text{Cor}(\mathbf{Z}_j, \mathbf{X}_k) = 0 \; \forall k$. These expectations are,

$$\text{E}[SSR_j] = \text{E}[\text{E}[SSR_j|\mathbf{X}]] = \sigma_y^2 + \sigma_c^2 + (n-2)\left(\sigma_j^2 + \sum_{i \neq j} r_{ij}^2 \sigma_i^2\right) \qquad (27)$$

$$\text{E}[SSE_j] = \text{E}[\text{E}[SSE_j|\mathbf{X}]] = (n-2)\left(\sigma_y^2 + \sigma_c^2 - \sigma_j^2 - \sum_{i \neq j} r_{ij}^2 \sigma_i^2\right) = (n-2)c_j(\sigma_y^2 + \sigma_c^2), \qquad (28)$$

where $c_j = 1 - \sum_i r_{ij}^2 h_{i,\mathrm{b}}^2$.

##### 3.4.2 Single-SNP analysis with covariates fitted, $\mathbf{Cor(X, Z) = 0}$.

We note that $\mathbf{Z\Sigma Z'} = \sigma_j^2 \mathbf{Z}_j \mathbf{Z}_j' + \sum_{i \neq j} \sigma_i^2 \mathbf{Z}_i \mathbf{Z}_i'$ and substitute the result into the RHS of (25), obtaining

$$
\begin{aligned}
&= \sigma_e^2 + \mathrm{Tr}((\mathbf{I} - \mathbf{X}(\mathbf{X}'\mathbf{X})^{-1}\mathbf{X}')\mathbf{Z}_j \mathbf{B}_j^{-1} \mathbf{Z}_j' (\mathbf{I} - \mathbf{X}(\mathbf{X}'\mathbf{X})^{-1}\mathbf{X}')\mathbf{Z\Sigma Z'}) \qquad (29)\\
&= \sigma_e^2 + \sum_{i \neq j} \mathrm{Tr}((\mathbf{I} - \mathbf{X}(\mathbf{X}'\mathbf{X})^{-1}\mathbf{X}')\mathbf{Z}_j \mathbf{B}_j^{-1} \mathbf{Z}_j' (\mathbf{I} - \mathbf{X}(\mathbf{X}'\mathbf{X})^{-1}\mathbf{X}')\mathbf{Z}_i \mathbf{Z}_i' \sigma_i^2) + \mathrm{Tr}((\mathbf{I} - \mathbf{X}(\mathbf{X}'\mathbf{X})^{-1}\mathbf{X}')\mathbf{Z}_j \mathbf{Z}_j' \sigma_j^2)
\end{aligned}
$$

For $\mathrm{Tr}((\mathbf{I} - \mathbf{X}(\mathbf{X}'\mathbf{X})^{-1}\mathbf{X}')\mathbf{Z}_j \mathbf{Z}_j' \sigma_j^2) = \mathbf{Z}_j' (\mathbf{I} - \mathbf{X}(\mathbf{X}'\mathbf{X})^{-1}\mathbf{X}')\mathbf{Z}_j \sigma_j^2 = \mathbf{B}_j \sigma_j^2$, we need

$$
\mathrm{E}[\mathbf{B}_j] = \mathrm{E}[\mathbf{Z}_j' (\mathbf{I} - \mathbf{X}(\mathbf{X}'\mathbf{X})^{-1}\mathbf{X}')\mathbf{Z}_j] = n{-}1 - \mathrm{E}[(n{-}1) \sum_{k=1}^{p} \hat{\mathrm{Cor}}(\mathbf{Z}_j, \mathbf{X}_k)^2] \approx n{-}1 - (n{-}1)\frac{p}{n{-}1} = n{-}1{-}p, \qquad (30)
$$

For the second trace in the expected value, rather than with $\mathbf{Z}_i$ directly, we use the equivalent

$$
\mathbf{Z}_i = \mathbf{Z}_j \hat{r}_{ij} + \hat{\mathbf{e}}_{ij},
$$

where $\hat{r}_{ij}$ is the sample correlation and the estimated coefficient for a simple linear regression of $\mathbf{Z}_i$ on $\mathbf{Z}_j$, since $\mathbf{Z}$ is standardised, and $\hat{\mathbf{e}}_{ij}$ is the estimated residual vector from the same regression,

$$
\begin{aligned}
&\mathrm{Tr}((\mathbf{I} - \mathbf{X}(\mathbf{X}'\mathbf{X})^{-1}\mathbf{X}')\mathbf{Z}_j \mathbf{B}_j^{-1} \mathbf{Z}_j' (\mathbf{I} - \mathbf{X}(\mathbf{X}'\mathbf{X})^{-1}\mathbf{X}')\mathbf{Z}_i \mathbf{Z}_i' \sigma_i^2)\\
&\quad = (\hat{r}_{ij} \mathbf{Z}_j' + \hat{\mathbf{e}}_{ij}')(\mathbf{I} - \mathbf{X}(\mathbf{X}'\mathbf{X})^{-1}\mathbf{X}')\mathbf{Z}_j \mathbf{B}_j^{-1} \mathbf{Z}_j' (\mathbf{I} - \mathbf{X}(\mathbf{X}'\mathbf{X})^{-1}\mathbf{X}')(\mathbf{Z}_j \hat{r}_{ij} + \hat{\mathbf{e}}_{ij})\sigma_i^2. \qquad (31)
\end{aligned}
$$

After expanding out terms and noting that by definition $\mathbf{e}_{ij}' \mathbf{Z}_j = 0$, (31) corresponds to

$$
\hat{r}_{ij}^2 \mathbf{B}_j \sigma_i^2 - 2\hat{r}_{ij} \hat{\mathbf{e}}_{ij}' \mathbf{X}(\mathbf{X}'\mathbf{X})^{-1}\mathbf{X}'\mathbf{Z}_j \sigma_i^2 + \hat{\mathbf{e}}_{ij}' \mathbf{X}(\mathbf{X}'\mathbf{X})^{-1}\mathbf{X}'\mathbf{Z}_j \mathbf{B}_j^{-1} \mathbf{Z}_j' \mathbf{X}(\mathbf{X}'\mathbf{X})^{-1}\mathbf{X}' \hat{\mathbf{e}}_{ij} \sigma_i^2. \qquad (32)
$$

We can write (32) in sample covariance notation, which due to standardisation is equivalent to correlation except for the case of $\hat{\mathrm{Cov}}(\hat{\mathbf{e}}_{ij}, \mathbf{X}_k) = \hat{\mathrm{Cor}}(\hat{\mathbf{e}}_{ij}, \mathbf{X}_k)\sqrt{1{-}\hat{r}_{ij}^2}$. Thus we have

$$
\begin{aligned}
&(n{-}1{-}p)\hat{r}_{ij}^2 \sigma_j^2 - 2(n{-}1)\hat{r}_{ij}\sqrt{1{-}\hat{r}_{ij}^2} \sum_{k=1}^{p} \hat{\mathrm{Cor}}(\hat{\mathbf{e}}_{ij}, \mathbf{X}_k)\hat{\mathrm{Cor}}(\mathbf{Z}_i, \mathbf{X}_k)\sigma_j^2 +\\
&\quad (1{-}\hat{r}_{ij}^2)(n{-}1) \sum_{k=1}^{p}\sum_{l=1}^{p}\sum_{m=1}^{p}\sum_{o=1}^{p} \frac{\hat{\mathrm{Cor}}(\hat{\mathbf{e}}_{ij}, \mathbf{X}_k)\hat{\mathrm{Cor}}(\mathbf{Z}_j, \mathbf{X}_k)\hat{\mathrm{Cor}}(\hat{\mathbf{e}}_{ij}, \mathbf{X}_l)\hat{\mathrm{Cor}}(\mathbf{Z}_j, \mathbf{X}_l)}{1{-}\hat{\mathrm{Cor}}(\mathbf{Z}_j, \mathbf{X}_m)\hat{\mathrm{Cor}}(\mathbf{Z}_j, \mathbf{X}_o)} \sigma_j^2. \qquad (33)
\end{aligned}
$$

To take the expectation of (33) with respect to $\mathbf{X}$, note if $k = l = m = o$, we have to scale the square of a $t$-distributed random variable $(n{-}2)\hat{\text{Cor}}(\mathbf{Z}_j, \mathbf{X}_k)^2/(1{-}\hat{\text{Cor}}(\mathbf{Z}_j, \mathbf{X}_k)^2)$, giving us an approximate expectation,

$$\left((n{-}1{-}p)\hat{r}_{ij}^2 + (1{-}\hat{r}_{ij}^2)\frac{np}{(n{-}2)^2}\right)\sigma_j^2, \tag{34}$$

which, marginalising over $\mathbf{Z}$ and assuming $n$ large, can be simplified:

$$\frac{n{-}p}{n{-}1}\left((n{-}2)r_{ij}^2 + 1\right)\sigma_j^2 + \frac{p}{n{-}2}\left(1 - \frac{(n{-}2)r_{ij}^2}{n{-}1} - \frac{1}{n{-}1}\right)\sigma_j^2. \tag{35}$$

The resulting approximation for $\text{E}[SSR_{j,2} - SSR_{j,1}]$ is,

$$\begin{aligned}
&\sigma_e^2 + (n{-}1{-}p)\sigma_j^2 + \left\{\frac{(n{-}1{-}p)(n{-}2) - p}{n{-}1}\right\}\sum_{i\neq j} r_{ij}^2\sigma_j^2 + \frac{(n{-}1{-}p) + p}{n{-}1}\sum_{i\neq j}\sigma_j^2 \\
&\quad = \sigma_e^2 + (n{-}1{-}p)\sigma_j^2 + (n{-}2{-}p)\sum_{i\neq j} r_{ij}^2\sigma_j^2 + \sum_{i\neq j}\sigma_j^2 \\
&\quad = \sigma_y^2 + (n{-}2{-}p)\sum_{i} r_{ij}^2\sigma_j^2.
\end{aligned} \tag{36}$$

For $\text{E}[SSE_{j,2}]$, we need the expectation of $\text{Tr}((\mathbf{I} - \mathbf{X}(\mathbf{X}'\mathbf{X})^{-1}\mathbf{X}')\mathbf{Z}\mathbf{\Sigma}\mathbf{Z}')$, which is,

$$\sum_{i=1}^{m}\text{E}[(n{-}1)\sigma_j^2 + (n{-}1)\sum_{k=1}^{p}\hat{\text{Cor}}(\mathbf{Z}_i, \mathbf{X}_k)^2\sigma_j^2] \approx \sum_{i=1}^{m}(n{-}1)\sigma_j^2 + (n{-}1)\frac{p}{n{-}1}\sigma_j^2 = (n{-}1{-}p)\sum_{i=1}^{m}\sigma_j^2 \tag{37}$$

Combined with (26) and (36), we obtain

$$\begin{aligned}
\text{E}[SSE_{j,2}] &\approx (n{-}2{-}p)\sigma_e^2 + (n{-}1{-}p)\sum_{i=1}^{m}\sigma_j^2 - (n{-}2{-}p)\sigma_j^2 - (n{-}2{-}p)\sum_{i\neq j} r_{ij}^2\sigma_j^2 - \sum_{i=1}^{m}\sigma_j^2 \\
&= (n{-}2{-}p)\left(\sigma_y^2 - \sigma_j^2 - \sum_{i\neq j} r_{ij}^2\sigma_j^2\right)
\end{aligned} \tag{38}$$

#### 3.5 Single-SNP analysis with covariates fitted, $\text{Cor}(\mathbf{X}, \mathbf{Z}) \neq \mathbf{0}$

In a multiple regression, the $t$ statistic for a covariate $i$ is conditional on all other covariates fitted in the model. Hence, statistics about $\mathbf{Z}_j$ from a regression with both $\mathbf{Z}_j$ and $\mathbf{X}$ fitted can be written as testing for an effect for the covariate $\hat{\mathbf{e}}_j$, where $\hat{\mathbf{e}}_j$ is the estimated residual from the regression,

$$\mathbf{Z}_j = \mathbf{X}\hat{\boldsymbol{\gamma}}_j + \hat{\mathbf{e}}_j. \tag{39}$$

Since $\mathbf{Z}$ is standardised, the estimated residual has sample variance $1-R_j^2$, where $R_j^2$ is the coefficient of determination, or equivalently the inverse of the variance inflation factor (Theil, 1971). Standardising $\hat{\mathbf{e}}_j$ to $\tilde{\mathbf{e}}_j$ we obtain to a linear model equivalent to (1):

$$\tilde{\mathbf{y}} = \mathbf{1}\mu + \tilde{\mathbf{e}}\boldsymbol{\alpha} + \boldsymbol{\varepsilon}, \quad \boldsymbol{\alpha} \sim \mathcal{N}(\mathbf{0}, \tilde{\boldsymbol{\Sigma}}), \boldsymbol{\varepsilon} \sim \mathcal{N}(\mathbf{0}, a\sigma_e^2\mathbf{I}), \tag{40}$$

where $\tilde{\boldsymbol{\Sigma}}_{jj} = (1-R_j^2)\sigma_j^2$, $\tilde{\mathbf{y}}$ is the covariate corrected phenotype and $a = (n-2-p)(n-2)$. We still need to convert the LD score from the observed to the residualised scale. To do this, we note

$$\begin{aligned}
\hat{r}_{ij} = \hat{\text{Cor}}(\mathbf{Z}_i, \mathbf{Z}_j) &:= \hat{\text{Cov}}(\mathbf{Z}_i, \mathbf{Z}_j) \\
&= \hat{\text{Cov}}(\mathbf{X}\hat{\boldsymbol{\gamma}}_i, \mathbf{X}\hat{\boldsymbol{\gamma}}_j) + \hat{\text{Cov}}(\mathbf{X}\hat{\boldsymbol{\gamma}}_i, \hat{\mathbf{e}}_j) + \hat{\text{Cov}}(\hat{\mathbf{e}}_i, \mathbf{X}\hat{\boldsymbol{\gamma}}_j) + \hat{\text{Cov}}(\hat{\mathbf{e}}_i, \hat{\mathbf{e}}_j) \\
&= \hat{\boldsymbol{\gamma}}_i' \text{Var}[\mathbf{X}]\hat{\boldsymbol{\gamma}}_j + \hat{\boldsymbol{\gamma}}_i' \hat{\text{Cov}}(\mathbf{X}, \hat{\mathbf{e}}_j) + \hat{\text{Cov}}(\hat{\mathbf{e}}_i, \mathbf{X})\hat{\boldsymbol{\gamma}}_j + \hat{\text{Cov}}(\hat{\mathbf{e}}_i, \hat{\mathbf{e}}_j) \\
&= \hat{\boldsymbol{\gamma}}_i' \hat{\text{Var}}[\mathbf{X}]\hat{\boldsymbol{\gamma}}_j + \hat{\text{Cor}}(\hat{\mathbf{e}}_i, \hat{\mathbf{e}}_j)\sqrt{(1-R_i^2)(1-R_j^2)},
\end{aligned} \tag{41}$$

giving an "LD score" on the residualised scale of,

$$\hat{r}'_{ij} = \text{Cor}(\hat{\mathbf{e}}_i, \hat{\mathbf{e}}_j) = \frac{\hat{r}_{ij} - \hat{\boldsymbol{\gamma}}_i' \hat{\text{Var}}[\mathbf{X}]\hat{\boldsymbol{\gamma}}_j}{\sqrt{(1-R_i^2)(1-R_j^2)}}. \tag{42}$$

substituting (40) and (42) into (8) and (10) respectively, with $\mathbf{X} = \mathbf{0}$, obtaining

$$\begin{aligned}
\text{E}[SSR_j|\mathbf{Z}, \mathbf{X}] &= a\sigma_e^2 + (n-1)(1-R_j^2)\sigma_j^2 + (n-1)\sum_{i \neq j}(\hat{r}'_{ij})^2(1-R_i^2)\sigma_i^2 \\
&= a\sigma_e^2 + (n-1)(1-R_j^2)\sigma_j^2 + (n-1)\sum_{i \neq j}\frac{(\hat{r}_{ij} - \hat{\boldsymbol{\gamma}}_i' \hat{\text{Var}}[\mathbf{X}]\hat{\boldsymbol{\gamma}}_j)^2(1-R_i^2)\sigma_i^2}{(1-R_i^2)(1-R_j^2)} \\
&= a\sigma_e^2 + (n-1)(1-R_j^2)\sigma_j^2 + (n-1)\sum_{i \neq j}\frac{(\hat{r}_{ij} - \hat{\boldsymbol{\gamma}}_i' \hat{\text{Var}}[\mathbf{X}]\hat{\boldsymbol{\gamma}}_j)^2}{(1-R_j^2)}\sigma_i^2.
\end{aligned} \tag{43}$$

For $SSE_j|\mathbf{Z}, \mathbf{X}$, the expectation can be determined by taking the expectations of the individual components, $SST|\mathbf{X} - SSR_j|\mathbf{Z}, \mathbf{X}$. These expected values are,

$$\begin{aligned}
\text{E}[SST|\mathbf{X}] &= (n-1)\left(a\sigma_e^2 + \sum_{i=1}^{m}(1-R_i^2)\sigma_i^2\right) \\
\text{E}[SSE_j|\mathbf{Z}, \mathbf{X}] &= (n-1)\left(a\sigma_e^2 + \sum_{i \neq j}(1-R_i^2)\sigma_i^2 - \sum_{i \neq j}\frac{(\hat{r}_{ij} - \hat{\boldsymbol{\gamma}}_i' \hat{\text{Var}}[\mathbf{X}]\hat{\boldsymbol{\gamma}}_j)^2}{(1-R_j^2)}\sigma_i^2\right) - a\sigma_e^2.
\end{aligned} \tag{44}$$

##### 3.6 Meta-analysis test statistics

We have shown that the definition of $h^2_{\text{SNP}}$ is dependent on the covariates fitted. Now, we show how this impacts estimation of $h^2_{\text{SNP}}$ from meta analysis summary statistics, including when there are individuals shared between studies. We focus on two types of meta-analysis, sample size weighted (Stouffer et al., 1949), where $T_{j,\text{Meta}}$ is calculated as,

$$\frac{\sum_s T_{j,s} w_s}{\sqrt{\sum_s w_s^2}}, \tag{45}$$

with $w_s = \sqrt{n_s}$, the sample size of study $s$, and inverse variance weighted (Willer et al., 2010), where a new estimate of the effect and the variance of the estimate is obtained and weights $w_s$ are equal to $1/\text{Var}[\hat{\alpha}_{j,s}]$,

$$\hat{\alpha}_{j,\text{Meta}} = \frac{\sum_s w_s \hat{\alpha}_{j,s}}{\sum_s w_s}, \quad \text{Var}[\hat{\alpha}_{j,\text{Meta}}] = \frac{1}{\sum_s w_s}. \tag{46}$$

Ideally, each study $s$ in the set $S$ would have performed identical analyses. In practice models may differ between studies. If the covariates missing from the analysis model in study $s$ are independent of the SNPs $\mathbf{Z}$, then unbiasedness of the estimated SNP effect $\hat{\alpha}_{j,s}$ would still hold for all $s$ (Kempthorne, 1977) if the response is continuous (Gail et al., 1984). This is sufficient for meta-analysis to gain statistical power at identifying causal loci without introducing bias, but insufficient for $h^2_{\text{SNP}}$ estimation. To prove this point, consider the case where there are always effects associated with $\mathbf{X}$, where $\mathbf{X}$ is standardised and independent of $\mathbf{Z}$ but $\mathbf{y}$ is not standardised. Now partition $S$ into the set $S_1$ where the analysis model ignored covariates and $S_2$ which did not,

$$\mathbf{y}_s = \mathbf{1}\mu + \mathbf{Z}_{j,s}\alpha_j + \boldsymbol{\varepsilon}, \qquad \text{if } s \in S_1 \tag{47}$$

$$\mathbf{y}_s = \mathbf{1}\mu + \mathbf{Z}_{j,s}\alpha_j + \mathbf{X}\boldsymbol{\beta} + \boldsymbol{\varepsilon} \qquad \text{if } s \in S_2. \tag{48}$$

For a sample-size weighted meta-analysis, and remembering that $\mathbf{X}\boldsymbol{\beta}$ is absorbed into $\boldsymbol{\epsilon}_s$ for studies $s \in S_1$,

$$\begin{aligned}
\text{E}[T^2_{j,\text{Meta}}] &= \frac{\text{E}_{\mathbf{Z}}[\text{E}_{\boldsymbol{\alpha},\boldsymbol{\varepsilon}}[(\sum_s T_{j,s} w_s)^2]]}{\sum_s w_s^2} \\
&= \frac{\text{E}_{\mathbf{Z}}[\text{E}_{\boldsymbol{\alpha},\boldsymbol{\varepsilon}}[(\sum_s \hat{\alpha}_{j,s} w_s / \sqrt{\text{Var}[\hat{\alpha}_{j,s})}]^2]]}{\sum_s w_s^2} \\
&\approx \frac{1}{\sum_s w_s^2} \text{E}_{\mathbf{Z}}\left[\text{E}_{\boldsymbol{\alpha},\boldsymbol{\varepsilon}}\left[\left(\sum_s \frac{\text{E}[\hat{\alpha}_{j,s}|\boldsymbol{\alpha},\boldsymbol{\varepsilon}] w_s}{\sqrt{\text{E}[\text{Var}[\hat{\alpha}_{j,s}]]}}\right)^2\right]\right]
\end{aligned}$$

$$
\begin{aligned}
&= \frac{1}{\sum_s w_s^2} \mathrm{E}_{\mathbf{Z}}\left[\mathrm{E}_{\boldsymbol{\alpha},\boldsymbol{\varepsilon}}\left[\left(\sum_s \frac{(\sum_i \hat{r}_{ij,s}\boldsymbol{\alpha}_i + c_s(\mathbf{Z}'_{j,s}\mathbf{Z}_{j,s})^{-1}\mathbf{Z}'_j\boldsymbol{\varepsilon}_s)w_s}{\sqrt{\mathrm{E}[\mathrm{Var}[\hat{\alpha}_{j,s}]]}}\right)^2\right]\right] \\
&= \frac{1}{\sum_s w_s^2}\left(\mathrm{E}_{\mathbf{Z}}\left[\sum_{s,s'} \frac{\mathrm{E}_{\boldsymbol{\alpha}}[(\sum_i \hat{r}_{ij,s}\boldsymbol{\alpha}_i)(\sum_i \hat{r}_{ij,s'}\boldsymbol{\alpha}_i)]w_s w_{s'}}{\sqrt{\mathrm{E}[\mathrm{Var}[\hat{\alpha}_{j,s}]]\mathrm{E}[\mathrm{Var}[\hat{\alpha}_{j,s'}]]}}\right] + \mathrm{E}_{\mathbf{Z}}\left[\mathrm{E}_{\boldsymbol{\varepsilon}}\left[\left(\sum_s \frac{c_s(\mathbf{Z}'_{j,s}\mathbf{Z}_{j,s})^{-1}\mathbf{Z}'_{j,s}\boldsymbol{\varepsilon}_s w_s}{\sqrt{\mathrm{E}[\mathrm{Var}[\hat{\alpha}_{j,s}]]}}\right)^2\right]\right]\right) \\
&= \frac{1}{\sum_s w_s^2}\left(\sum_i \sigma_i^2 \sum_{s,s'} \frac{\hat{r}_{ij,s}\hat{r}_{ij,s'}w_s w_{s'}}{\sqrt{\mathrm{E}[\mathrm{Var}[\hat{\alpha}_{j,s}]]\mathrm{E}[\mathrm{Var}[\hat{\alpha}_{j,s'}]]}} + \sum_{s,s'} \frac{(\sigma_e^2 + \mathbb{1}_{s,s'\in S_1}\sigma_c^2)c_s c_{s'} w_s w_{s'}(n_{ss'}-1)}{(n_s-1)(n_{s'}-1)\sqrt{\mathrm{E}[\mathrm{Var}[\hat{\alpha}_{j,s}]]\mathrm{E}[\mathrm{Var}[\hat{\alpha}_{j,s'}]]}}\right),
\end{aligned}
\tag{49}
$$

where $c_s = 1$ and $c_s = \sqrt{(n_s-1)/(n_s-p-1)}$ for analysis models (47) and (48). To complete the derivation of the approximate expectation, we need

$$
\mathrm{E}[\mathrm{Var}[\hat{\alpha}_{j,s}]] = \begin{cases} \frac{\sigma_y^2+\sigma_c^2-\sum_i r_{ij}^2\sigma_i^2}{n_s-1} \approx \frac{\sigma_y^2+\sigma_c^2}{n_s-1} \approx \frac{\sigma_y^2+\sigma_c^2}{w_s^2} & \text{if } c_s = 1 \\ \frac{\sigma_y^2-\sum_i r_{ij}^2\sigma_i^2}{n_s-p-1} \approx \frac{\sigma_y^2}{n_s-p-1} \approx \frac{\sigma_y^2}{c_s^2 w_s^2} & \text{if } c_s = \sqrt{(n_s-p-1)/(n_s-1)}. \end{cases}
\tag{50}
$$

If $c_s = 1 \forall s$ or $c_s = \sqrt{(n_s-p-1)/(n_s-1)}\forall s$ and we assume $p \ll n_s \forall s \Rightarrow c_s \approx 1$, (49) reduces to,

$$
\begin{aligned}
\mathrm{E}[T_{j,\mathrm{Meta}}^2] &= \frac{1}{\sum_s w_s^2}\left(\sum_i \sigma_i^2 \sum_{s,s'} \frac{\hat{r}_{ij,s}\hat{r}_{ij,s'}c_s c'_s w_s^2 w_{s'}^2}{\sigma_T^2} + (\sigma_T^2 - \sum_i \sigma_i^2)\sum_{s,s'} \frac{c_s^2 c_{s'}^2 w_s^2 w_{s'}^2(n_{ss'}-1)}{(n_s-1)(n'_s-1)\sigma_T^2}\right), \\
&\approx \frac{1}{\sum_s w_s^2}\left(\sum_i \sigma_i^2 \sum_{s,s'} \frac{\hat{r}_{ij,s}\hat{r}_{ij,s'}w_s^2 w_{s'}^2}{\sigma_T^2} + (\sigma_T^2 - \sum_i \sigma_i^2)\sum_{s,s'} \frac{n_{ss'}}{\sigma_T^2}\right),
\end{aligned}
\tag{51}
$$

where $\sigma_T^2 = \sigma_y^2 + \sigma_c^2$ and noting that $\sigma_e^2 + \mathbb{1}_{s,s'\in S_1}\sigma_c^2 = \sigma_T^2 - \sum_i \sigma_i^2$ when no covariate correction was applied, and $\sigma_y^2$ if covariate correction was applied. When the approximate expectation for the sample LD in studies $s$ and $s'$,

$$
\mathrm{E}[\hat{r}_{ij,s}\hat{r}_{ij,s'}] = r_{ij}^2 + \frac{n_{ss'}}{n_s n_{s'}}
\tag{52}
$$

is substituted into (51), we find

$$
\begin{aligned}
\mathrm{E}[T_{j,\mathrm{Meta}}^2] &\approx \frac{1}{\sum_s w_s^2}\left(\sum_i \frac{r_{ij}^2\sigma_i^2}{\sigma_T^2}\sum_s w_s^2 \sum_{s'} w_{s'}^2 + \frac{\sigma_T^2 - \sum_i \sigma_i^2 + \sum_i \sigma_i^2}{\sigma_T^2}\sum_{s,s'} n_{ss'}\right), \\
&= \sum_s n_s \sum_i \frac{r_{ij}^2\sigma_i^2}{\sigma_T^2} + \frac{\sum_{s,s'} n_{ss'}}{\sum n_s}, \\
&\approx \begin{cases} 1 + \frac{2}{n}\sum_{s\neq s'} n_{ss'} + n\sum_i r_{ij}^2 h_{i,\mathrm{b}}^2 & \text{if } c_s = 1 \quad \forall s \\ 1 + \frac{2}{n}\sum_{s\neq s'} n_{ss'} + n\sum_i r_{ij}^2 h_{i,\mathrm{a}}^2 & \text{if } c_s = \sqrt{(n_s-1)/(n_s-p-1)} \quad \forall s \end{cases}.
\end{aligned}
\tag{53}
$$

Unlike the single study considered earlier, $\mathrm{E}[T^2_{j,\mathrm{Meta}}]$ can be inflated in such a way that the inclusion of an intercept term is sufficient to obtain unbiased estimates of $h^2_{\mathrm{SNP}}$. However this inflation is due to individuals being included in multiple studies, not the explicit ignoring of confounders we focus on in this paper. If studies from sets $S_1$ and $S_2$, are combined, a meaningful interpretation of the slope as a function of a heritability parameter is no longer possible, as in

$$\begin{aligned}
\mathrm{E}[T^2_{j,\mathrm{Meta}}] &= \frac{\sum_i \sigma_i^2}{\sum_s w_s^2}\Bigg(\sum_{s,s'\in S_1}\frac{\sqrt{(n_s-1)(n_{s'}-1)}\hat{r}_{ij,s}\hat{r}_{ij,s'}w_s w_{s'}}{\sigma_y^2+\sigma_c^2}+\sum_{s,s'\in S_2}\frac{\sqrt{(n_s-p-1)(n_{s'}-p-1)}\hat{r}_{ij,s}\hat{r}_{ij,s'}w_s w_{s'}}{\sigma_y^2} \\
&\quad + \sum_{s\in S_1,s'\in S_2}\frac{\sqrt{(n_s-1)(n_{s'}-p-1)}w_s w_{s'}\hat{r}_{ij,s}\hat{r}_{ij,s'}}{\sigma_y^2\sqrt{1+\sigma_c^2/\sigma_y^2}}\Bigg) \\
&\quad + \frac{\sigma_e^2}{\sum_s w_s^2}\Bigg(\sum_{s,s'\in S_2}\frac{w_s w_s'}{\sigma_y^2}+\sum_{s\in S_1,s'\in S_2}\frac{w_s w_s'}{\sigma_y^2\sqrt{1+\sigma_c^2/\sigma_y^2}}\Bigg)+\frac{\sigma_e^2+\sigma_c^2}{\sum_s w_s^2}\sum_{s,s'\in S_1}\frac{w_s w_s'}{\sigma_y^2+\sigma_c^2}, \qquad (54)
\end{aligned}$$

the denominator changes depending on the study's choice of $S_1$ or $S_2$. Similar problems arise in the case of inverse variance weighting, both for the test statistic $T^2_{j,\mathrm{Meta}}$ and the effect size $\hat{\alpha}_{j,\mathrm{Meta}}$. The expected value of $\hat{\alpha}^2_{j,\mathrm{Meta}}$ from an inverse variance weighted meta-analysis is,

$$\begin{aligned}
\mathrm{E}[\hat{\alpha}^2_{j,\mathrm{Meta}}] &= \mathrm{E}[(\frac{\sum_s \hat{\alpha}_{j,s}w_s}{\sum_s w_s})^2] \approx \frac{\sum_{s,s'}\mathrm{E}[(\hat{\alpha}_{j,s}\hat{\alpha}_{j,s'}]w_s w_{s'}}{(\sum_s w_s)^2} \\
&= \sum_i \sigma_i^2\frac{\sum_{s,s'}(r_{ij}^2+\frac{n_{ss'}}{n_s n_{s'}})w_s w_{s'}}{(\sum_s w_s)^2}+\sum_{s,s'}(\sigma_e^2+\mathbb{1}_{s,s'\in S_2}\sigma_c^2)\frac{n_{ss'}c_s c_{s'}w_s w_{s'}}{(n_s-1)(n_{s'}-1)(\sum_s w_s)^2} \\
&\approx \sum_i r_{ij}^2\sigma_i^2\frac{(\sum_s w_s)^2}{(\sum_s w_s)^2}+\sum_{s,s'}(\sigma_e^2+\mathbb{1}_{s,s'\in S_2}\sigma_c^2)\frac{n_{ss'}c_s c_{s'}w_s w_{s'}}{n_s n_{s'}(\sum_s w_s)^2}+\sum_i\sigma_i^2\sum_{s,s'}\frac{n_{ss'}w_s w_{s'}}{n_s n_{s'}(\sum_s w_s)^2} \\
&= \sum_i r_{ij}^2\sigma_i^2+\sum_{s,s'}(\sigma_e^2+\mathbb{1}_{s,s'\in S_1}\sigma_c^2)\frac{n_{ss'}c_s c_{s'}w_s w_{s'}}{n_s n_{s'}(\sum_s w_s)^2}+\sum_i\sigma_i^2\sum_{s,s'}\frac{n_{ss'}w_s w_{s'}}{n_s n_{s'}(\sum_s w_s)^2}, \qquad (55)
\end{aligned}$$

The expected values of the weights in (55) are,

$$w_s \approx 1/\mathrm{E}[\mathrm{Var}[\hat{\alpha}_{j,s}]] = \frac{n_s-1}{\sigma_y^2+\sigma_c^2-\sum_i r_{ij}^2\sigma_i^2} \approx \frac{n_s-1}{\sigma_y^2+\sigma_c^2}, \quad \text{if } c_s = 1 \qquad (56)$$

$$w_s \approx 1/\mathrm{E}[\mathrm{Var}[\hat{\alpha}_{j,s}]] = \frac{n_s-p-1}{\sigma_y^2-\sum_i r_{ij}^2\sigma_i^2} \approx \frac{n_s-p-1}{\sigma_y^2} = \frac{c_s^2(n_s-1)}{\sigma_y^2} \text{ if } c_s = \sqrt{\frac{n_s-p-1}{n_s-1}}. \qquad (57)$$

which upon substitution into (55) and again assuming $p \ll n$ so that $c_s \approx 1\,\forall s$ gives,

$$\mathrm{E}[\hat{\alpha}^2_{j,\mathrm{Meta}}] \approx \begin{cases} \left(\frac{1}{\sum_s n_s - 1} + \frac{2\sum_{s\neq s'} n_{ss'}}{\left(\sum_s n_s - 1\right)^2}\right)(\sigma_y^2 + \sigma_c^2) + \sum_i r_{ij}^2 \sigma_i^2 & \text{if } c_s = 1 \quad \forall s \\ \left(\frac{1}{\sum_s n_s - 1} + \frac{2\sum_{s\neq s'} n_{ss'}}{\left(\sum_s n_s - 1\right)^2}\right)\sigma_y^2 + \sum_i r_{ij}^2 \sigma_i^2 & \text{if } c_s = \sqrt{\frac{n_s - 1}{n_s - p - 1}} \quad \forall s \\ \frac{(\sigma_y^2 + \sigma_c^2)\sum_{s,s' \in S_1} n_{ss'}}{\left(\sum_s n_s - 1\right)^2} + \frac{\sigma_y^2 \sum_{s,s' \in S_2} n_{ss'}}{\left(\sum_s n_s - 1\right)^2} + \frac{\sigma_y^2 \sum_{s \in S_1, s' \in S_2} n_{ss'}}{\left(\sum_s n_s - 1\right)^2} + \sum_i r_{ij}^2 \sigma_i^2 & \text{in general.} \end{cases} ,(58)$$

Similarly, the test statistic $T^2_{j,\mathrm{Meta}} = \hat{\alpha}^2_{j,\mathrm{Meta}} \sum_s w_s$ has an approximate expected value of

$$\mathrm{E}[T^2_{j,\mathrm{Meta}}] \approx \begin{cases} 1 + \frac{2\sum_{s\neq s'} n_{ss'}}{\sum_s n_s} + \sum_s n_s \sum_i r_{ij}^2 h_{i,\mathrm{b}}^2 & \text{if } c_s = 1 \forall s \\ 1 + \frac{2\sum_{s\neq s'} n_{ss'}}{\sum_s n_s - 1} + \sum_s n_s \sum_i r_{ij}^2 h_{i,\mathrm{a}}^2 & \text{if } c_s = \sqrt{\frac{n_s - 1}{n_s - p - 1}} \forall s \\ \left(\frac{(\sigma_y^2 + \sigma_c^2) \sum\limits_{s,s' \in S_1} n_{ss'} + \sigma_y^2 (\sum\limits_{s,s' \in S_2} n_{ss'} + \sum\limits_{s \in S_1, s' \in S_2} n_{ss'})}{\left(\sum\limits_s n_s\right)^2} + \sum\limits_i r_{ij}^2 \sigma_i^2\right)(\sum\limits_{s \in S_1} \frac{n_s - 1}{\sigma_y^2 + \sigma_c^2} + \sum\limits_{s \in S_2} \frac{n_s - p - 1}{\sigma_y^2}) & \text{in general.} \end{cases} ,(59)$$

so like sample weighted meta-analysis shows additive inflation in the presence of shared individuals, and thus a meaningful heritability parameter cannot be estimated unless different studies apply equivalent covariate corrections. The presence of additive inflation should be viewed as analogous to the expectation of the cross product of test statistics given in Bulik-Sullivan et al. (2015b) in order to estimate genetic correlation.

In the main text we showed that if Cov[$\mathbf{X}, \mathbf{Z}$]=0, the SumHer model intercept is 1 and there is no need to fit $A/C$, including in meta-analysis with equivalent treatment of covariates. In Figure 5 of the main text (for LDAK simulations) and in Figure S9 (for GCTA simulations), we show what happens if we repeat this analysis with sample overlap. Specifically the meta-analysis now consists of two studies, one with all 8000 individuals and the with 4000 of the individuals. Based on the theory given in section 3.6, either $h^2_{\mathrm{SNPa}}$ or $h^2_{\mathrm{SNPb}}$ can still be estimated without/with covariate correction in both studies, but with model intercept $2 \times 4000/12000 = 5/3$. Our simulations are consistent with this theory under both LDAK and GCTA models.

#### 3.7 Mixed model test statistics

##### 3.7.1 Determining the expectation

For the purposes of this section, we still assume the same phenotype model as in (1) where $\mu$ is an intercept, $\mathbf{Z}$ a $n \times m$ matrix of standardised SNP genotypes, $\boldsymbol{\alpha}$ a vector of SNP effect sizes, and $\mathbf{\Sigma}$ a diagonal matrix with $j$th entry $\sigma_j^2$. However, the model fitted in the association study now is,

$$\mathbf{y} = \mathbf{1}\mu + \mathbf{Z}_j\boldsymbol{\alpha}_j + \mathbf{Z}\boldsymbol{\alpha}_{-j} + \boldsymbol{\varepsilon}, \quad \boldsymbol{\alpha}_{-j} \sim \mathcal{N}(\mathbf{0}, \tilde{\mathbf{\Sigma}}), \boldsymbol{\varepsilon} \sim \mathcal{N}(\mathbf{0}, \tilde{\sigma}_e^2\mathbf{I}), \tag{60}$$

where the effect of SNP $j$ is treated as a fixed effect, and the elements of $\tilde{\mathbf{\Sigma}}$ and $\tilde{\sigma}_e^2$ need not match the equivalent entries in the true model. The test statistic for testing the hypothesis whether $\boldsymbol{\alpha}_j = 0$ calculated under the assumed model of (60) is,

$$T^2_{j,LMM} = \mathbf{y}'\mathbf{V}^{-1}\mathbf{Z}_j(\mathbf{Z}_j'\mathbf{V}^{-1}\mathbf{Z}_j)^{-1}\mathbf{Z}_j'\mathbf{V}^{-1}\mathbf{y} \tag{61}$$

where $\mathbf{V} = \mathbf{Z}_{-j}\tilde{\mathbf{\Sigma}}\mathbf{Z}_{-j}' + \tilde{\sigma}_e^2\mathbf{I}$. Note that part of the simplicity of (61) is due to the centring required to standardise $\mathbf{Z}$, which implies $\text{cov}(\hat{\mu}, \hat{\alpha}_j) = 0$. Now

$$\begin{aligned}
\text{E}[T^2_{j,LMM}] &= \text{E}[\mathbf{y}'\mathbf{V}^{-1}\mathbf{Z}_j(\mathbf{Z}_j'\mathbf{V}^{-1}\mathbf{Z}_j)^{-1}\mathbf{Z}_j'\mathbf{V}^{-1}\mathbf{y}] \\
&= Tr(\mathbf{V}^{-1}\mathbf{Z}_j(\mathbf{Z}_j'\mathbf{V}^{-1}\mathbf{Z}_j)^{-1}\mathbf{Z}_j'\mathbf{V}^{-1}\text{E}[\mathbf{y}\mathbf{y}']) \\
&= Tr(\mathbf{V}^{-1}\mathbf{Z}_j(\mathbf{Z}_j'\mathbf{V}^{-1}\mathbf{Z}_j)^{-1}\mathbf{Z}_j'\mathbf{V}^{-1}(\mu^2\mathbf{1}\mathbf{1}' + \sigma_j^2\mathbf{Z}_j\mathbf{Z}_j' + \mathbf{Z}_{-j}\mathbf{\Sigma}_{-j}\mathbf{Z}_{-j}' + \sigma_e^2\mathbf{I})) \\
&= Tr(\mathbf{V}^{-1}\mathbf{Z}_j(\mathbf{Z}_j'\mathbf{V}^{-1}\mathbf{Z}_j)^{-1}\mathbf{Z}_j'\mathbf{V}^{-1}(\sigma_j^2\mathbf{Z}_j\mathbf{Z}_j' + \mathbf{Z}_{-j}\mathbf{\Sigma}_{-j}\mathbf{Z}_{-j}' + \sigma_e^2\mathbf{I})) \\
&= \mathbf{Z}_j'\mathbf{V}^{-1}\mathbf{Z}_j(\mathbf{Z}_j'\mathbf{V}^{-1}\mathbf{Z}_j)^{-1}\mathbf{Z}_j'\mathbf{V}^{-1}\mathbf{Z}_j\sigma_j^2 + Tr(\mathbf{V}^{-1}\mathbf{Z}_j(\mathbf{Z}_j'\mathbf{V}^{-1}\mathbf{Z}_j)^{-1}\mathbf{Z}_j'\mathbf{V}^{-1}(\mathbf{Z}_{-j}\mathbf{\Sigma}_{-j}\mathbf{Z}_{-j}' + \sigma_e^2\mathbf{I})) \\
&= \mathbf{Z}_j'\mathbf{V}^{-1}\mathbf{Z}_j\sigma_j^2 + Tr(\mathbf{V}^{-1}\mathbf{Z}_j(\mathbf{Z}_j'\mathbf{V}^{-1}\mathbf{Z}_j)^{-1}\mathbf{Z}_j'\mathbf{V}^{-1}(\mathbf{Z}_{-j}\mathbf{\Sigma}_{-j}\mathbf{Z}_{-j}' + \sigma_e^2\mathbf{I})). \qquad (62)
\end{aligned}$$

Lastly, define $\mathbf{V}_{\text{True}} = \mathbf{Z}_{-j}\mathbf{\Sigma}_{-j}\mathbf{Z}_{-j}' + \sigma_e^2\mathbf{I}$ and $\mathbf{E} = \mathbf{V}_{\text{True}} - \mathbf{V}$. Then we can write (62) as,

$$\begin{aligned}
\text{E}[T^2_{j,LMM}] &= \mathbf{Z}_j'\mathbf{V}^{-1}\mathbf{Z}_j\sigma_j^2 + Tr(\mathbf{V}^{-1}\mathbf{Z}_j(\mathbf{Z}_j'\mathbf{V}^{-1}\mathbf{Z}_j)^{-1}\mathbf{Z}_j'\mathbf{V}^{-1}(\mathbf{V} + \mathbf{E})) \\
&= \mathbf{Z}_j'\mathbf{V}^{-1}\mathbf{Z}_j\sigma_j^2 + 1 + \mathbf{Z}_j'\mathbf{V}^{-1}\mathbf{E}\mathbf{V}^{-1}\mathbf{Z}_j(\mathbf{Z}_j'\mathbf{V}^{-1}\mathbf{Z}_j)^{-1}. \qquad (63)
\end{aligned}$$

For the purposes of trying to re-estimate $h^2_{\text{SNP}}$ from mixed model association statistics, while $\mathbf{Z}_j'\mathbf{V}^{-1}\mathbf{Z}_j = \text{Var}[\hat{\boldsymbol{\beta}}]$ is a readily obtainable quantity, $\mathbf{Z}_j'\mathbf{V}^{-1}$ is not. More importantly, re-estimation of $h^2_{\text{SNP}}$ would only be of interest in the case $\mathbf{E} \neq \mathbf{0}$.

##### 3.7.2 Pitfalls of approximating $\text{E}[T^2_{j,LMM}]$ as a linear function of $\text{E}[T^2_{j,LMM}]$

In practice many users will be tempted to assume $\text{E}[T^2_{j,LMM}]$ is similar in form to $\text{E}[T^2_{j,LM}]$,

$$\text{E}[T^2_{j,LMM}] \approx a + \sum_i r_{ij}^2 h_j^2, \tag{64}$$

Since (63) implies this is not true, a more accurate view is

$$\mathrm{E}[T_{j,LMM}^2] \approx a + b\mathrm{E}[T_{j,LM}^2], \tag{65}$$

where $\mathrm{E}[T_{j,LM}^2]$ is known from results in sections 3.1-3.6. Substituting the form of $\mathrm{E}[T_{j,LM}^2]$ into (65),

$$\mathrm{E}[T_{j,LMM}^2] \approx a + b(1 + \sum_i r_{ij}^2 h_j^2), \tag{66}$$

we find that, unless $a = 0$ and b is unconstrained, or a is unconstrained and $b = 1$, only biased estimates of $h^2_{\mathrm{SNP}}$ can be estimated, corresponding to either $bh^2_{\mathrm{SNP}}$ (assuming confounding is accounted by A) or $bh^2_{\mathrm{SNP}}/(a+b)$ (assuming confounding is accounted by C).

### 4 Additional simulation results

#### 4.1 Phenotypes generated assuming under a GCTA model

To show that our conclusions are not dependent on the simulation model, we simulated phenotypes using a GCTA model and analysed using LDSC. Results are presented in Figures S1-S4.

In Figure S1(a-c), we see the same patterns as in the LDAK simulations, namely ignoring covariates means only $h^2_{\mathrm{SNPb}}$ can be estimated, while conditioning on covariates, means only $h^2_{\mathrm{SNPa}}$ can be estimated from $T_j^2$, while both $h^2_{\mathrm{SNPa}}$ and $h^2_{\mathrm{SNPb}}$ can be estimated from $n\hat{\alpha}_j^2$. By changing the subset of SNPs, the degree of negative bias when estimating $h^2_{\mathrm{SNP}}$ from $n\hat{\alpha}_j^2$ and fixing the intercept to one when covariates were adjusted for is less than for the LDAK simulations presented in the main paper.

For meta-analysis test statistics (Figure S2), we find as expected the same patterns as in the LDAK simulations. If there is consistency in the GWAS analysis method, a meaningful definition of $h^2_{\mathrm{SNP}}$ is estimated. If there is inconsistency in analysis models between component GWAS $s$, the estimate of $h^2_{\mathrm{SNP}}$ obtained from a meta-analysis is not interpretable.

In Figure S3, we see only full adjustment for confounders in the original GWAS leads to unbiased estimation of $h^2_{\mathrm{SNPa}}$. As with the LDAK simulations, biased $h^2_{\mathrm{SNP}}$ estimates were obtained in both the no adjustment and partial adjustment case at the GWAS stage. Compared to the LDAK simulations, the level of bias in $h^2_{\mathrm{SNP}}$ was lower when simulating under a GCTA model (Figures S3, S4).

Like the LDAK simulations, we find that the positive bias in the estimate of $h^2_{\mathrm{SNPb}}$ and the scale of $A/C$ (Figure S4) for $C1$, $C2$, and $C3$ phenotypes correspond to the proportion of phenotypic variation attributed to confounding. Again like the LDAK simulations, and as expected both from the positive linear relationship between LD score and $a_j$ and from results in DeVlaming et al. (2017), the estimates of $A/C$ were below the mean level of confounding.

#### 4.2 Estimation of $h^2_{\text{SNP}}$ when applying filtering

In many of the simulations with unaccounted confounding, we generate $S_j$ with values sufficiently high that SNP $j$ would be removed from analysis using the filtering methods proposed in Zheng et al. (2017) as an outlier. We applied this filtering approach to see what impact it had on the estimate of $h^2_{\text{SNP}}$ and $A/C$ from phenotypes simulated under both a GCTA and LDAK model.

The LDAK simulation results presented in Figures S5-S6 are equivalent to Figures 3-4 in the main paper, but with filtering applied before summary statistic $h^2_{\text{SNP}}$ analysis. The filtering consisted of removing SNPs where $S_j > 80$ and also those SNPs in linkage disequilibrium (defined as $r^2_{ij} > 0.10$) with such SNPs from analysis. Similarly the GCTA simulations results presented in Figures S7-S8 are equivalent to Figures S3-S4, but with filtering applied before summary statistic $h^2_{\text{SNP}}$ analysis.

While as expected, the estimates of $h^2_{\text{SNP}}$ drop compared to the no filtering case for both LDAK and GCTA simulations, in no case was the drop sufficient to eliminate bias. While some of the GCTA simulations may appear to be unbiased, with the average estimate of $h^2_{\text{SNP}}$ sitting on the black line, it is always the red line that is the estimable definition of $h^2_{\text{SNP}}$ in these simulations. Furthermore, as the filtering is based on a fixed cut-off, as the proportion of phenotypic variation, $\sigma^2_c/(\sigma^2_y + \sigma^2_c)$ due to confounding increased, an increasing number of SNPs were removed in the filtering process. This meant the patterns linking $\sigma^2_c/(\sigma^2_y + \sigma^2_c)$ to the level of bias in the estimate of $h^2_{\text{SNP}}$ are broken as seen in Figures S6 and S8. From a statistical perspective, this indicates ad-hoc filtering approaches may be insufficient to remove all influential SNPs that could bias the estimation of parameters.

#### 4.3 Comparing heritability models

While not of primary interest, for the sake of completeness we have chosen to demonstrate how the choice of model for $h^2_j$ can adversely impact estimation. To do this, we compare the component of the expected test statistic due to polygenicity, $n\sum_i r^2_{ij}h^2_j$. For this, we consider four models. The first two are already proposed in the literature, GCTA (Yang et al., 2011), and LDAK (Speed et al., 2012). The other two are somewhat artificial cases designed to illustrate the impact of concentrating causal variants into specific regions defined by either minor allele frequency and LD. In the minor allele frequency scenario, we assume only variants $j$ where MAF $< 0.03$ are causal, which corresponds to 61,719 out of the 558,431 SNPs available. In the LD scenario, we assume only variants $j$ where $\sum_i r^2_{ij} > 13$ are causal, which corresponds to 59,311 out of the 558,431 SNPs available. For both the MAF and high LD scenario, we assume $h^2_j$ for causal variants is equal for all $j$. The results are presented in Figure 6 of the main paper.

Jie Zheng, A. Mezut Erzurumluoglu, Benjamin L. Elsworth, John P. Kemp, Laurence Howe, Philip C. Haycock, Gibran Hemani, Katherine Tansey, Charles Laurin, Early Genetics and Lifecourse Epidemiology (EAGLE) Consortium, Beate St Pourcain, Nicole M. Warrington, Hilary K. Finucane, Alkes L. Price, B. K. Bulik-Sullivan, Verneri Anttila, Lavinia Paternoster, Tom R. Gaunt, David M. Evans, and Benjamin M. Neale. LD Hub: a centralized database and web interface to perform LD score regression that maximizes the potential of summary level GWAS data for SNP heritability and genetic correlation analysis. *Bioinformatics*, 33(2):272–279, 2017.

J. Yang, S. Hong Lee, M.E. Goddard, and P. M. Visscher. Gcta: A tool for genome-wide complex trait analysis. *The American Journal of Human Genetics*, 88:76–82, 2011.

D. Speed, G. Hemani, M. R. Johnson, and D. J. Balding. Improved heritability estimation from genome-wide SNPs. *The American Journal of Human Genetics*, 91:1011–1021, 2012.

### 5 Supporting Figures

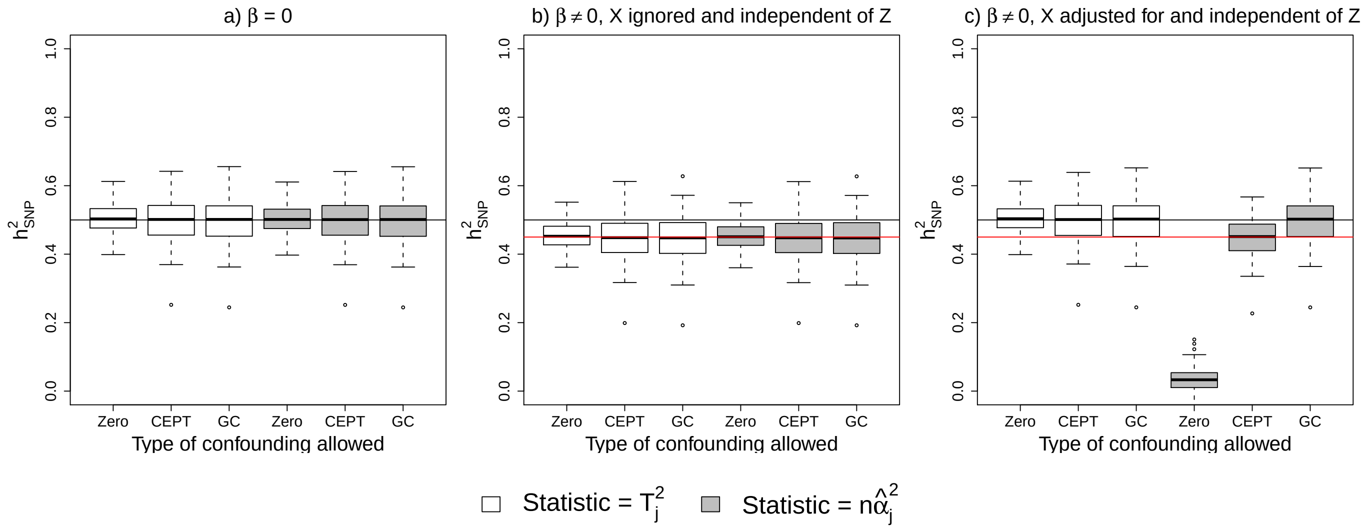


Figure S1: Estimates of $h^2_{\mathrm{SNP}}$ obtained from LDSC analysis of GWAS summary statistics constructed for phenotypes simulated under a GCTA model. The black and red horizontal lines indicate the values of $h^2_{\mathrm{SNPa}}$ and $h^2_{\mathrm{SNPb}}$. Zero, CEPT and GC refer to no, $A$ and $C$ confounding terms in the analysis model. (a) Phenotypes with no covariate effects. (b) Phenotypes with covariate effects but $\mathbf{X}$ ignored in the analysis. (c) Phenotypes with covariate effects and $\mathbf{X}$ adjusted for in the analysis.

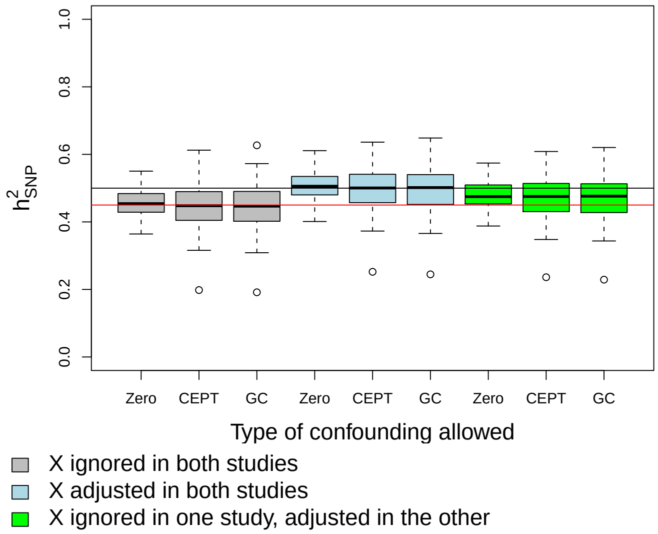


Figure S2: Estimates of $h^2_{\text{SNP}}$ obtained from LDSC analysis of summary statistics calculated from a meta-analysis of two GWAS constructed from phenotypes simulated under a GCTA model.

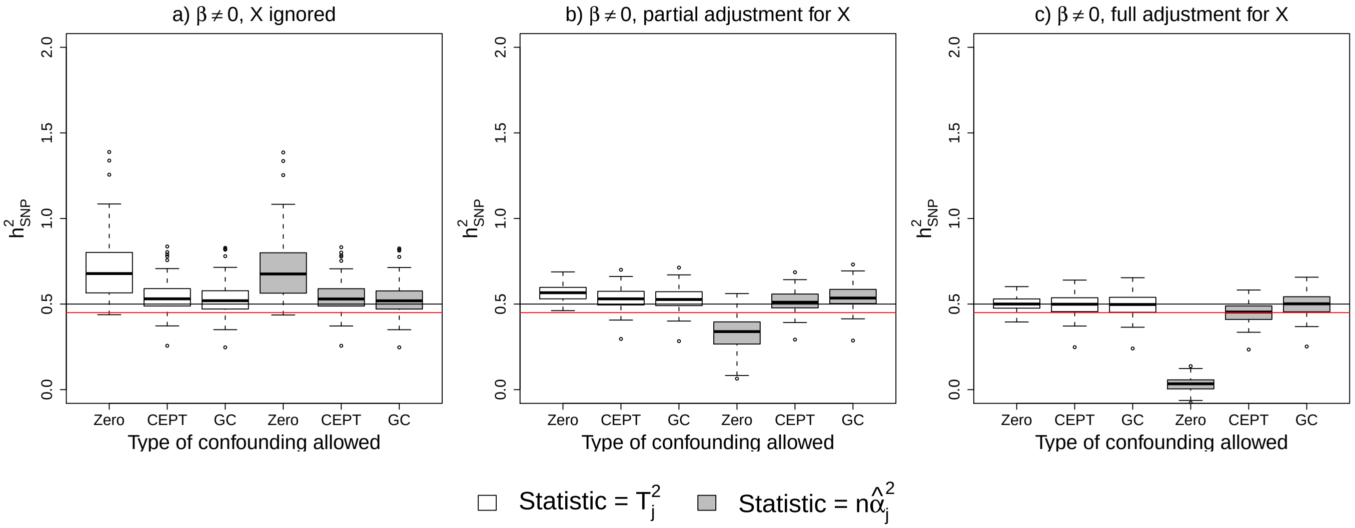


Figure S3: Similar to Figure S1, but here GWAS phenotypes (specifically C1) are subject to confounding: phenotype means differ among three subpopulations that each consist of three sub-subpopulations. The sub-populations were the same as used in the LDAK simulations. Estimates of $h^2_{\text{SNP}}$ from a GWAS with (a) no covariate adjustment, (b) adjustment for the three subpopulations but not the sub-subpopulations, (c) full covariate adjustment.

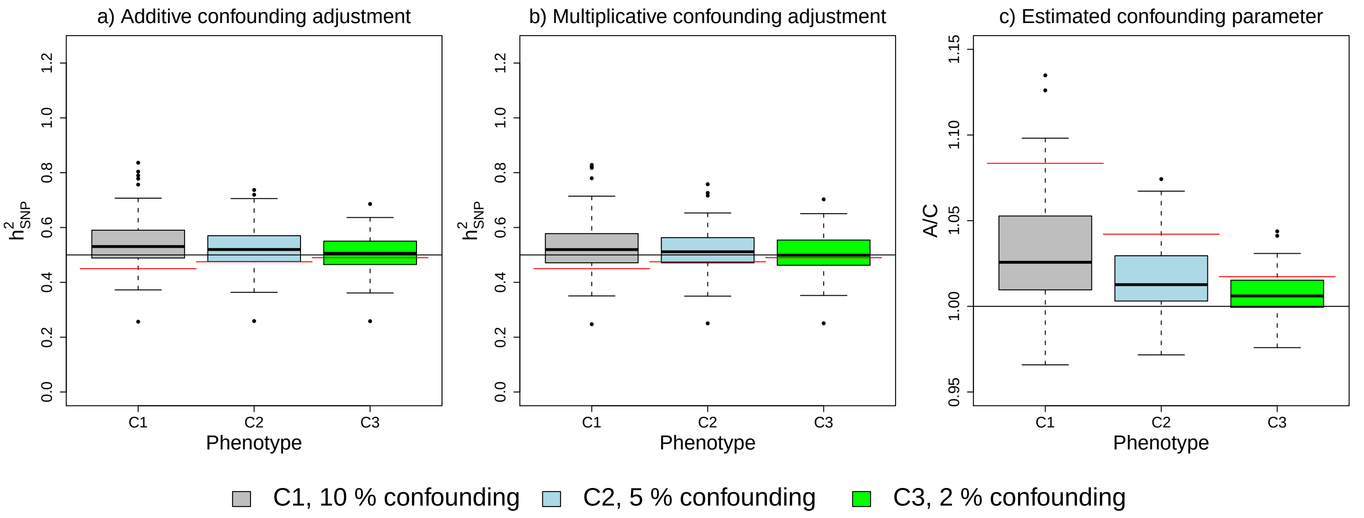


Figure S4: Estimating $h^2_{\text{SNP}}$ and confounding parameters from phenotypes with differing proportion of phenotypic variation (C1: 10 %, C2: 5 %, C3: 2 %) due to confounding when $h^2_{\text{SNPa}} = 0.5$. The confounding corresponds to ignoring sub-populations, which are the same as used in the LDAK simulations. The black lines in (a,b) indicates the simulated value of $h^2_{\text{SNPa}}$ and the red lines the simulated value of $h^2_{\text{SNPb}}$. In (c), the black line corresponds to $A/C = 1$, which is normally assumed to indicate no confounding and the red line the mean level of confounding, estimated as $\bar{S}_j - 1 - n\overline{\sum_i r^2_{ij} h^2_{j,\text{b}}}$. Note that the $y$-axis differs between plots.

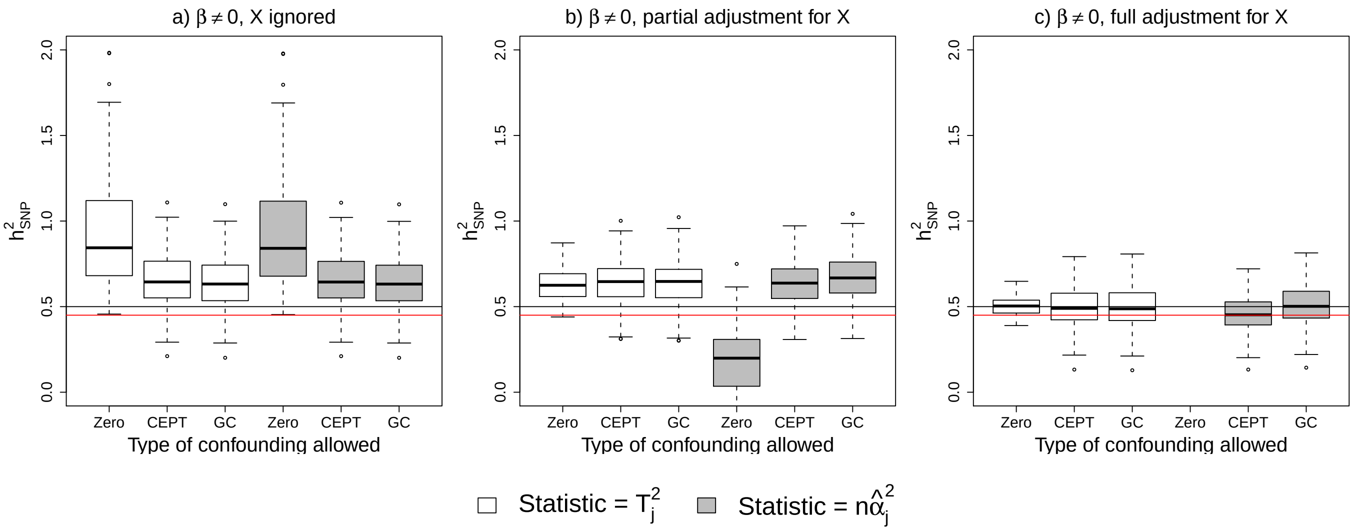


Figure S5: Same as Figures 3 in the main paper, but with filtering out of SNPs where $S_j > 80$ prior to summary statistic $h^2_{\text{SNP}}$ analysis. Estimates of $h^2_{\text{SNP}}$ from a GWAS with (a) no covariate adjustment, (b) adjustment for the three subpopulations but not the sub-subpopulations, (c) full covariate adjustment.

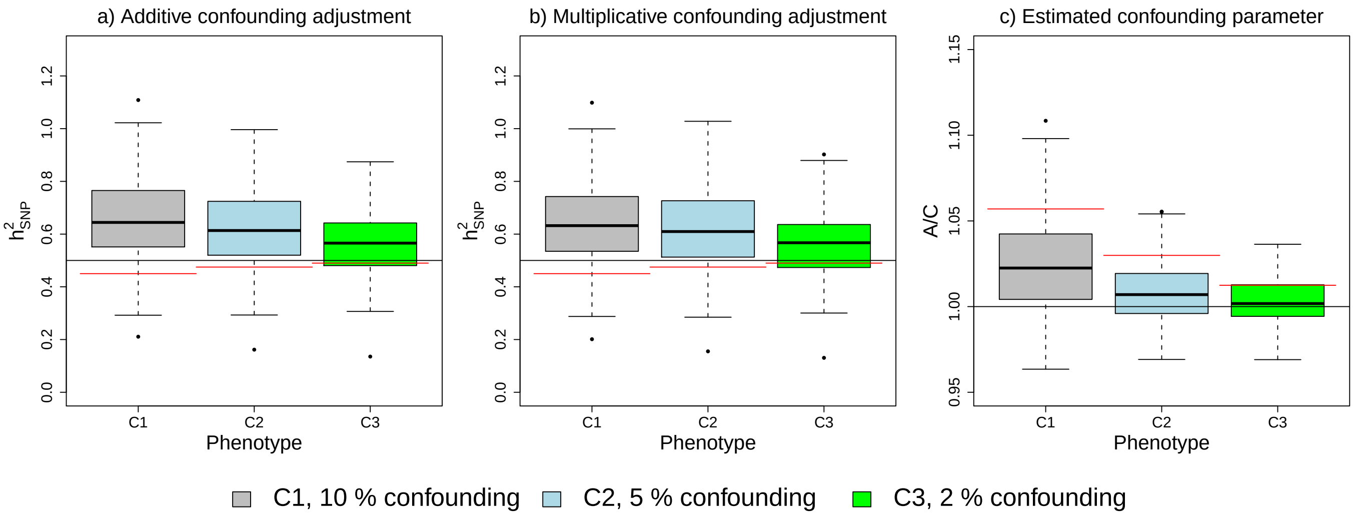


Figure S6: Same as Figure 4 in main paper, but with filtering out of SNPs where $S_j > 80$ prior to summary statistic $h^2_{\mathrm{SNP}}$ analysis. The black lines in (a,b) indicates the simulated value of $h^2_{\mathrm{SNPa}}$ and the red lines the simulated value of $h^2_{\mathrm{SNPb}}$. In (c), the black line corresponds to $A/C = 1$, which is normally assumed to indicate no confounding and the red line the mean level of confounding, estimated as $\bar{S}_j - 1 - n\overline{\sum_i r^2_{ij} h^2_{j,\mathrm{b}}}$. Note that the $y$-axis differs between plots.

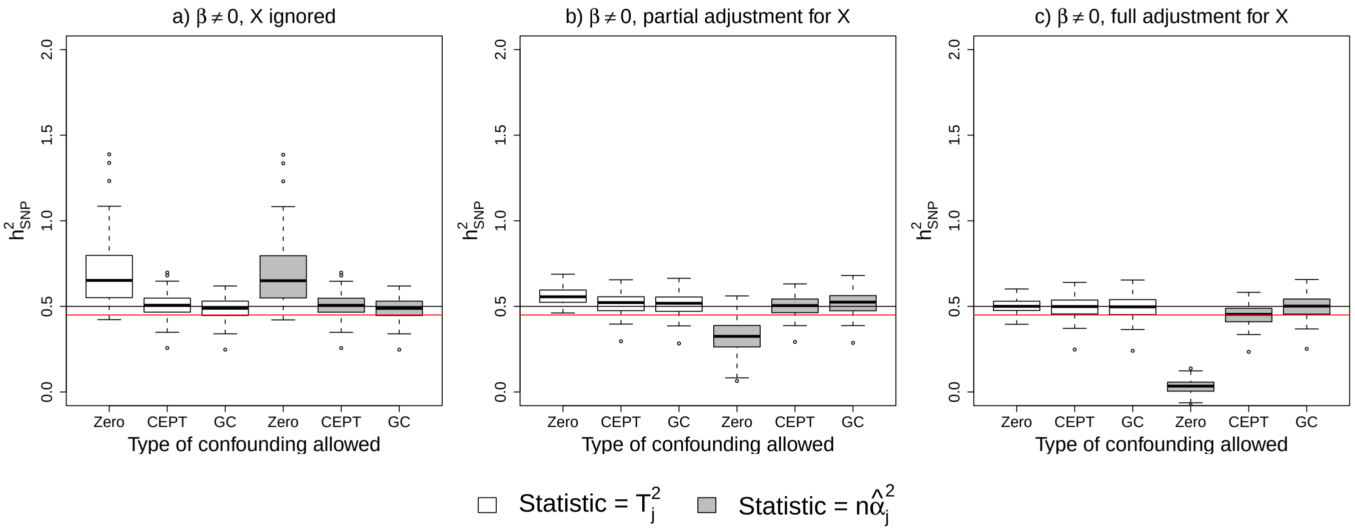


Figure S7: Same as Figure S3, but with filtering out of SNPs where $S_j > 80$ prior to summary statistic $h^2_{\mathrm{SNP}}$ analysis. Estimates of $h^2_{\mathrm{SNP}}$ from a GWAS with (a) no covariate adjustment, (b) adjustment for the three subpopulations but not the subsubpopulations, (c) full covariate adjustment.

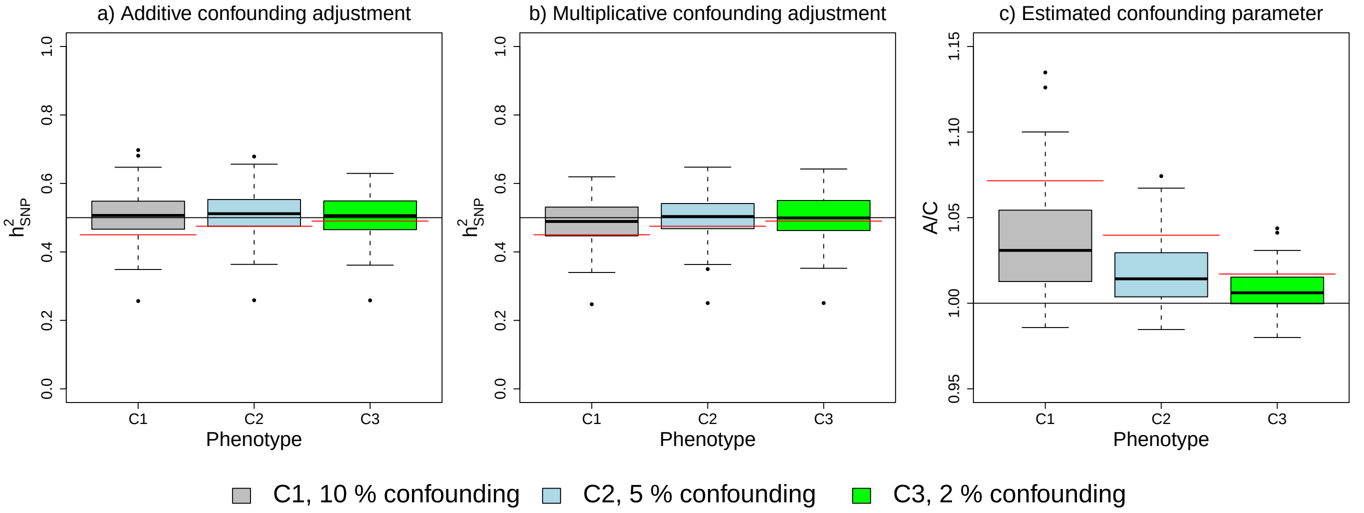


Figure S8: Same as Figure S4, but with filtering out of SNPs where $S_j > 80$ prior to summary statistic $h^2_{\mathrm{SNP}}$ analysis. The black lines in (a,b) indicates the simulated value of $h^2_{\mathrm{SNPa}}$ and the red lines the simulated value of $h^2_{\mathrm{SNPb}}$. In (c), the black line corresponds to $A/C = 1$, which is normally assumed to indicate no confounding and the red line the mean level of confounding, estimated as $\bar{S}_j - 1 - n\overline{\sum_i r^2_{ij} h^2_{j,\mathrm{b}}}$. Note that the $y$-axis differs between (a,b) and (c).

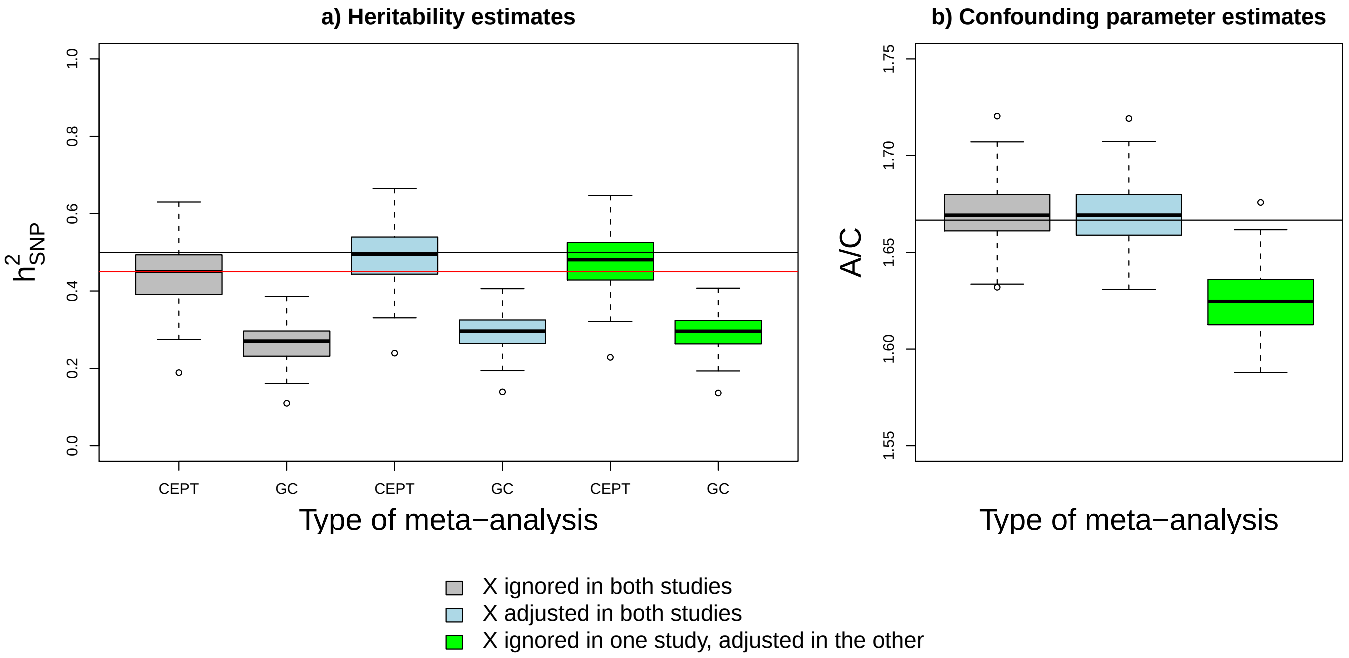


Figure S9: Same as Figure 5 in the main paper, with the meta-analysis including 4000 overlapping individuals out of a total of 12,000. The black and red horizontal lines in (a,c) indicate the values of $h^2_{\mathrm{SNPa}}$ and $h^2_{\mathrm{SNPb}}$. The black horizontal line in (b,d) indicates the expected intercept. CEPT and GC refer to A and C confounding terms in the analysis. Plot (a) gives estimates of $h^2_{\mathrm{SNP}}$ obtained under either an A or C view of the confounding parameter, while plot (b) gives estimates of the confounding parameter. Phenotypes were simulated in accordance to a GCTA model.

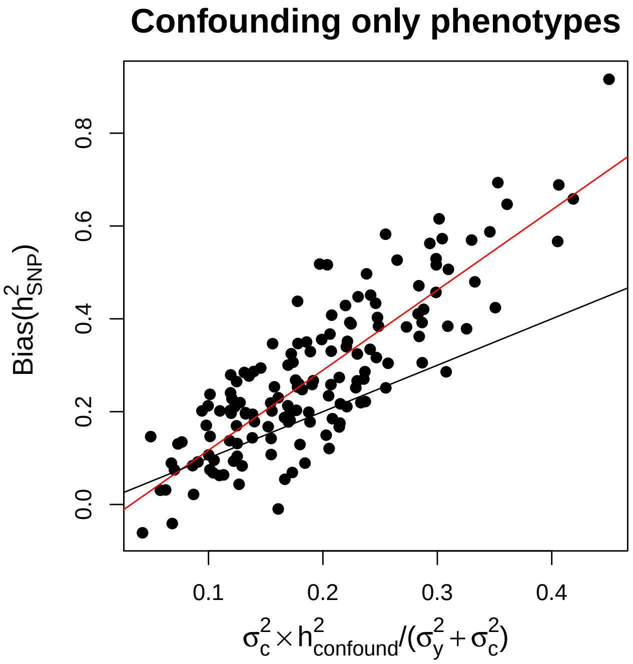


Figure S10: Estimates of $h^2_{\mathrm{SNP}}$ (called $h^2_{\mathrm{confound}}$) when the phenotype is the confounding effect $\mathbf{X}\boldsymbol{\beta}$ only, scaled to match the contribution of confounding to a C1 phenotype compared to the observed bias in $h^2_{\mathrm{SNP}}$ when confounding correction was not and was included in the GWAS. The $y = x$ line is shown in black, which corresponds to the bias in $h^2_{\mathrm{SNP}}$ being completely explained by the confounding-only test statistic. The red line is the fit of simple linear regression of the bias on scaled $h^2_{\mathrm{confound}}$. Bias was calculated for each phenotype using $\hat{h}^2_{\mathrm{SNP,ignored}} - \hat{h}^2_{\mathrm{SNP,corrected}}(h^2_{\mathrm{SNPb}}/h^2_{\mathrm{SNPa}})$, i.e by comparing estimates from a confounding ignored and a confounding corrected GWAS while accounting for the change in estimable $h^2_{\mathrm{SNP}}$ parameter. One confounding-only phenotypes did not produce convergent estimates so was removed for comparison purposes.
